# Specificities of and functional coordination between the two Cas6 maturation endonucleases in *Anabaena* sp. PCC 7120 assign orphan CRISPR arrays to three groups

**DOI:** 10.1101/2020.04.14.041012

**Authors:** Viktoria Reimann, Marcus Ziemann, Hui Li, Tao Zhu, Juliane Behler, Xuefeng Lu, Wolfgang R. Hess

**Author notes:** Vetter Pharma Fertigung GmbH & Co. KG, Schützenstraße 87, D-88212 Ravensburg, Germany.

## Abstract

The majority of bacteria and archaea possess an RNA-guided adaptive and inheritable immune system against viruses and other foreign genetic elements that consists of clustered regularly interspaced short palindromic repeats (CRISPRs) and CRISPR-associated (Cas) proteins. In most CRISPR-Cas systems, the maturation of CRISPR-derived small RNAs (crRNAs) is essential for functionality. In some bacteria, multiple instances of *cas* gene-free (orphan) repeat-spacer arrays exist, while additional instances of arrays that are linked to *cas* gene cassettes are present elsewhere in the genome.

In the cyanobacterium *Anabaena* sp. PCC 7120, ten CRISPR-Cas repeat-spacer arrays are present, but only two *cas* gene cassettes plus a Tn7-associated eleventh array are observed. In this study, we deleted the two *cas6* genes *alr1482* (Type III-D) or *alr1566* (Type I-D) and tested the specificities of the two corresponding enzymes in the resulting mutant strains, as recombinant proteins and in a cell-free transcription-translation system. The results assign the direct repeats (DRs) to three different groups. While Alr1566 is specific for one group, Alr1482 has a higher preference for the DRs of the second group but can also cleave those of the first group. We found that this cross-recognition limits crRNA accumulation for the Type I-D system *in vivo*.

We also show that the DR of the *cas* gene-free CRISPR array of cyanophage N-1 is processed by these enzymes, suggesting that it is fully competent in association with host-encoded Cas proteins. The data support a strong tendency for array fragmentation in multicellular cyanobacteria and disfavor other possibilities, such as the nonfunctionality of these orphan repeat-spacer arrays. Our data demonstrate the functional coordination of Cas6 endonucleases with both neighboring and remote repeat-spacer arrays in the CRISPR-Cas system of cyanobacteria.

## Introduction

Clustered regularly interspaced short palindromic repeats (CRISPR) and CRISPR-associated (Cas) proteins provide an adaptive immune system directed against invading nucleic acids in many bacteria and most archaea (Barrangou et al., 2007; Bolotin et al., 2005; Jansen et al., 2002; Makarova et al., 2006; Mojica et al., 2005, 2009; Pourcel et al., 2005). Immunity relies on the transcription of the CRISPR repeat-spacer array, ultimately leading to short CRISPR RNAs (crRNAs) that guide the CRISPR complexes in the recognition and destruction of invading nucleic acids. In most systems, first, a long precursor transcript (pre-crRNA) originates from the repeat-spacer array (Brouns et al., 2008; Hale et al., 2008) that is cleaved into short crRNAs (Hochstrasser and Doudna, 2015). The maturation of crRNAs is essential for the functionality of CRISPR-Cas systems and facilitates the recognition and destruction of invading nucleic acids upon base pairing through the interference complex (Charpentier et al., 2015; Hochstrasser and Doudna, 2015). In Type I, III, V and VI systems, specialized endoribonucleases recognize their cognate direct repeat (DR) sequences and mediate crRNA maturation (Behler and Hess, 2019; Hale et al., 2008, 2009; Karginov and Hannon, 2010; Przybilski et al., 2011). In the majority of Type I and III systems, the cognate endoribonucleases belong to the Cas6 protein family (Carte et al., 2008; Hale et al., 2008; Hochstrasser and Doudna, 2015). However, these specialized endoribonucleases can also be replaced by host enzymes in some instances (Behler et al., 2018; Chou-Zheng and Hatoum-Aslan, 2019).

Cyanobacteria are of considerable interest in biotechnology and ecology, as they are the only bacteria that perform oxygenic photosynthesis. In addition to the fixation of inorganic carbon through photosynthesis, some species are additionally capable of converting atmospheric nitrogen into organic biomass, thereby sustaining diazotrophic growth. Some of these cyanobacteria are multicellular and feature complex genomes. In these strains, the presence of multiple orphan CRISPR repeat-spacer cassettes not linked to any *cas* genes is typical (Hou et al., 2019). Free-standing repeat-spacer arrays were also observed in up to ~11% of previously investigated CRISPR systems of other phyla, although these findings might include some false-CRISPR elements (Makarova et al., 2015; Zhang and Ye, 2017). Among the multicellular, filamentous cyanobacteria, the high frequency of orphan CRISPR arrays is illustrated by such examples as *Tolypothrix bouteillei* VB521301 with 44 repeat-spacer arrays but only five *cas1* genes or the strain *Nostoc calcicola* FACHB-389, which has 19 instances of repeat-spacer arrays and only two *cas1* genes (Hou et al., 2019).

The strain *Anabaena* (*Nostoc*) sp. PCC 7120 (from here: *Anabaena* 7120) is a model for the group of multicellular, nitrogen-fixing cyanobacteria. A previously performed search for interspaced DR sequences suggested the existence of 11 CRISPR-like repeat-spacer cassettes in *Anabaena* 7120, which were designated CR_1 to CR_11 (Hou et al., 2019). One of these cassettes, CR_9, belongs to a Tn7-associated CRISPR derivative similar to those described in *Scytonema hofmanni* UTEX B 2349 and *Anabaena cylindrica* PCC 7122, two other multicellular cyanobacteria (Strecker et al., 2019).

All 11 arrays are transcribed and showed the crRNA-typical accumulation patterns of precursors and matured crRNAs, which is in contrast to the fact that only two instances of *cas* gene cassettes were found in this organism plus one additional isolated CRISPR-Cas integration cassette containing a *cas1, cas2* and a *csx18* gene (Hou et al., 2019; Shah et al., 2019). According to the specific *cas* gene content found in these cassettes, the existence of a Type I-D and a III-D system in *Anabaena* 7120 was predicted (Hou et al., 2019). These *cas* gene cassettes are linked to CR_4 (the I-D system) or CR_2 and CR_3 in the case of the III-D system. Hence, together with the Tn7-linked CR_9 array, four arrays appear connected to a particular system, while the other seven (CR_1, CR_5 to CR_8 and CR_10 and CR_11) qualify as orphan arrays. In addition, a short CRISPR array was described in cyanophage N-1, for which *Anabaena* 7120 is a suitable host (Adolph and Haselkorn, 1971, 1973). This CRISPR array was demonstrated to be transcribed in *Anabaena* 7120 during infection (Chénard et al., 2016).

In this study, we deleted the two *cas6* genes in *Anabaena* 7120 and tested the specificities of the two corresponding enzymes in the resulting mutant strains as recombinant proteins and in a cell-free transcription-translation (TXTL) system.

The results assign five different arrays each to one or to the other of the two Cas6 riboendonucleases and clearly show that processing of the Tn7-associated CR_9 is independent from either of the two enzymes. Hence, the data support a strong tendency for array fragmentation in multicellular cyanobacteria and disfavor other possibilities, such as that these orphan repeat-spacer arrays are nonfunctional. In addition, we observed a striking *in vivo* cross-specificity between the two Cas6 enzymes in which the presence of one protein lowers the processing and crRNA accumulation of maturation products produced by the other enzyme.

## Materials and methods

### Cultures and construction of mutant strains

Cultures of *Anabaena* 7120 were grown photoautotrophically in BG11 liquid medium or on agar plates (Rippka et al., 1979) under white light illumination of 30-50 μmol photons m^-2^ s^-1^ at 30 °C. The medium was supplemented with 30 μg mL^-1^ neomycin if needed.

The CRISPR-Cas12a (Cpf1) genome editing tool together with the pSL2680 plasmid (Addgene No. 85581) was used for the construction of *Anabaena* mutants, as previously reported (Ungerer and Pakrasi, 2016). First, a pair of complementary oligonucleotides that target the genes to be knocked out were annealed, treated with T4 polynucleotide kinase (Thermo Fisher), and cloned into the *AarI* (Thermo Fisher) digested plasmid pSL2680 by T4 DNA ligase (NEB), resulting in the gRNA-cassette editing plasmids. Second, the repairing templates (1 kb upstream and downstream flanked regions, respectively) were amplified from genomic DNA of *Anabaena* 7120 and inserted into the *KpnI* (Thermo Fisher) sites of the editing plasmids by seamless assembly, resulting in the final gRNA and repairing cassette editing plasmids. Third, the final editing plasmids were introduced into wild-type *Anabaena* 7120 by triparental mating conjugal transfer. Finally, genotypes of exconjugants were confirmed by PCR (**Figure S1**), and the editing plasmids were diluted and finally removed by subculture in BG11 medium without antibiotics. Primer pairs of *alr1482gRNA-1/2* and *alr1566gRNA-1/2* were used to prepare the gRNA-cassette editing plasmids, and primer pairs of *alr1482KO-1/2* and *alr1566*KO-1/2 were used to prepare the gRNA and repairing cassette editing plasmids for constructing *Δalr1482* and *Δalr1566*, respectively. Conjugal transfer was performed as previously reported (Elhai and Wolk, 1988). The plasmids pRL443 and pRL623 and the final CRISPR editing plasmids generated in this study were used as conjugal, helper and cargo plasmids, respectively. The sequences of all oligonucleotides are listed in **Table 1**. All PCR fragments, plasmids generated in this study, and gene mutation regions in the mutants were verified by Sanger sequencing.

**Table 1.**
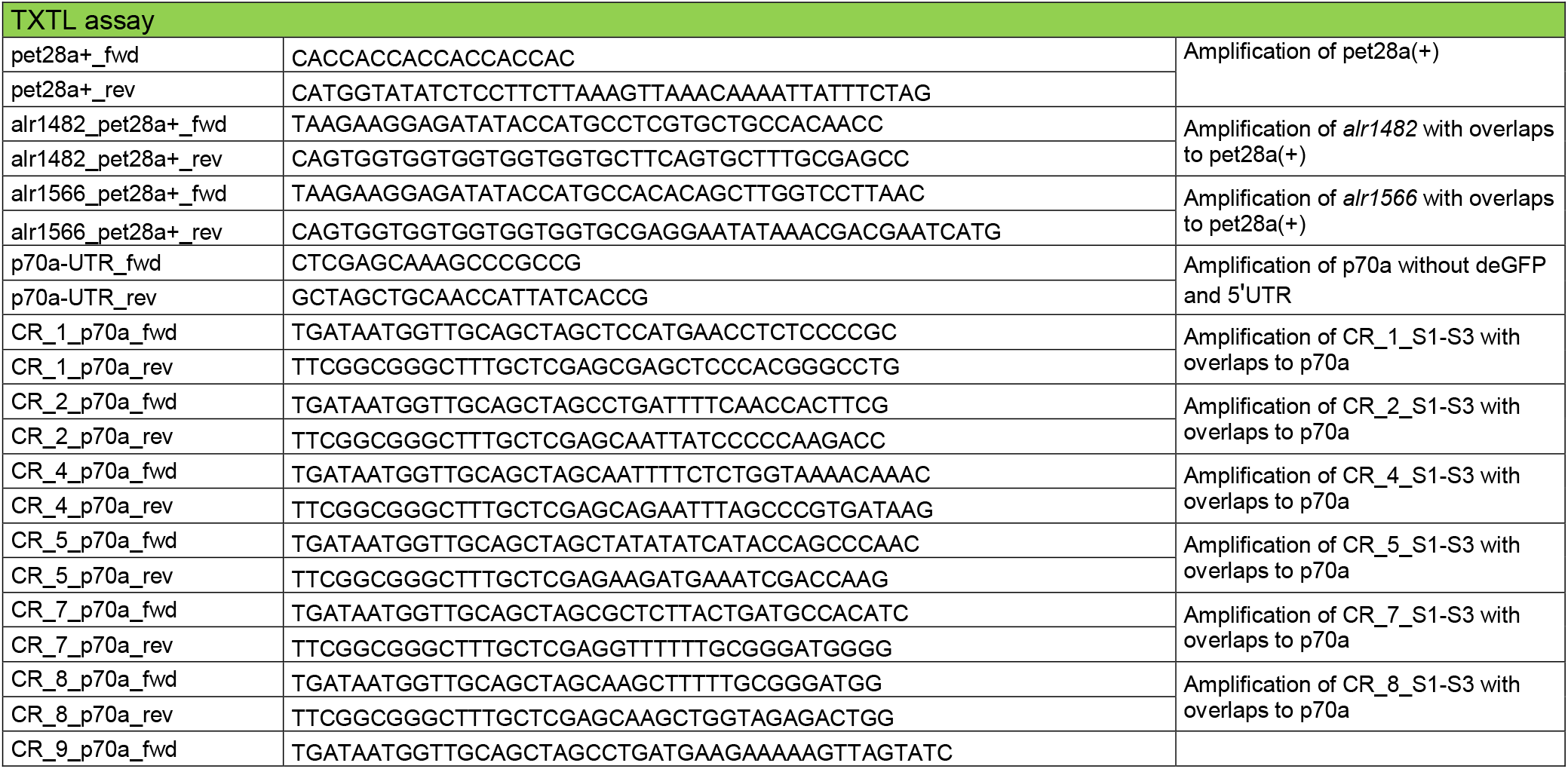
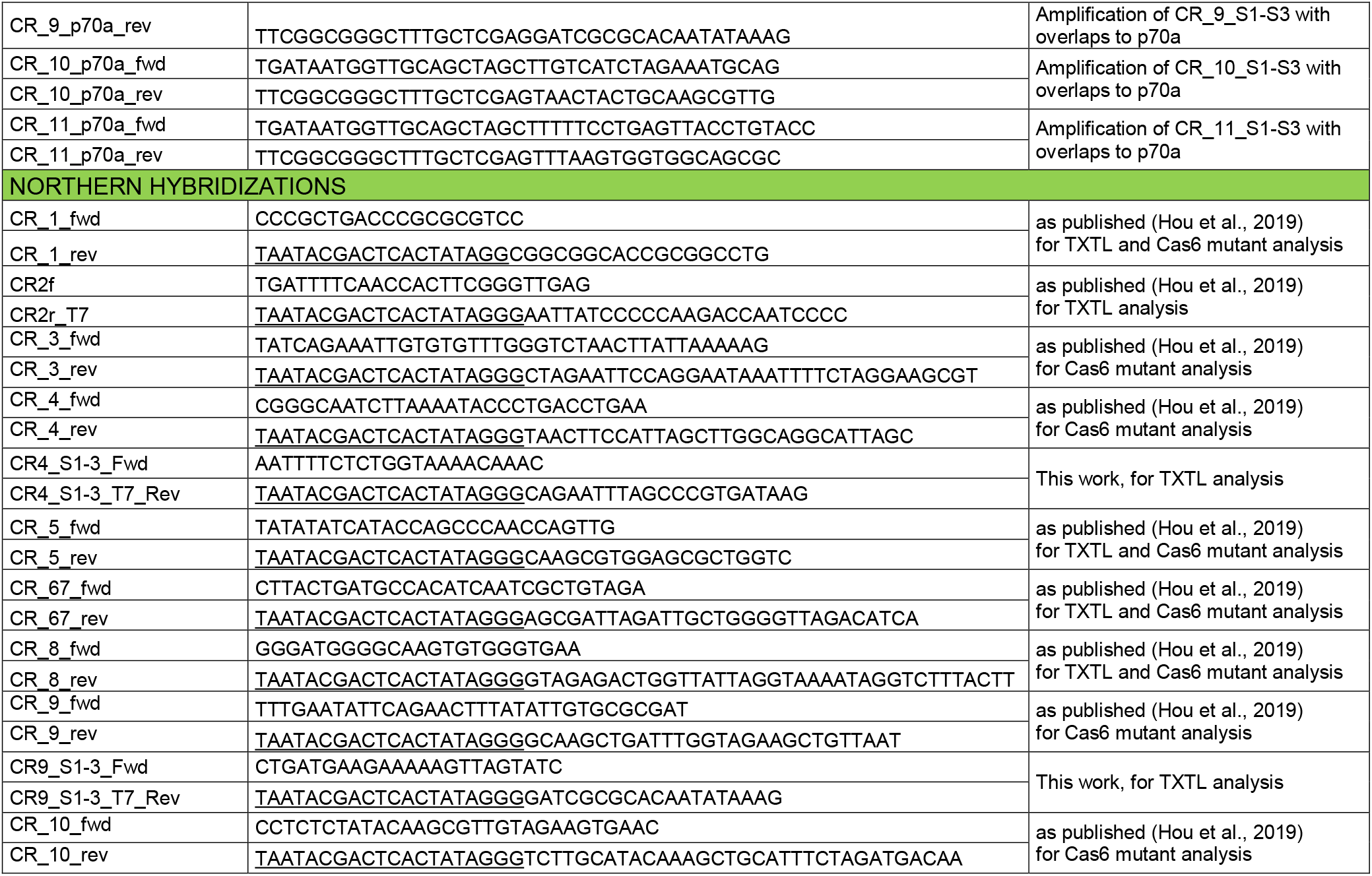
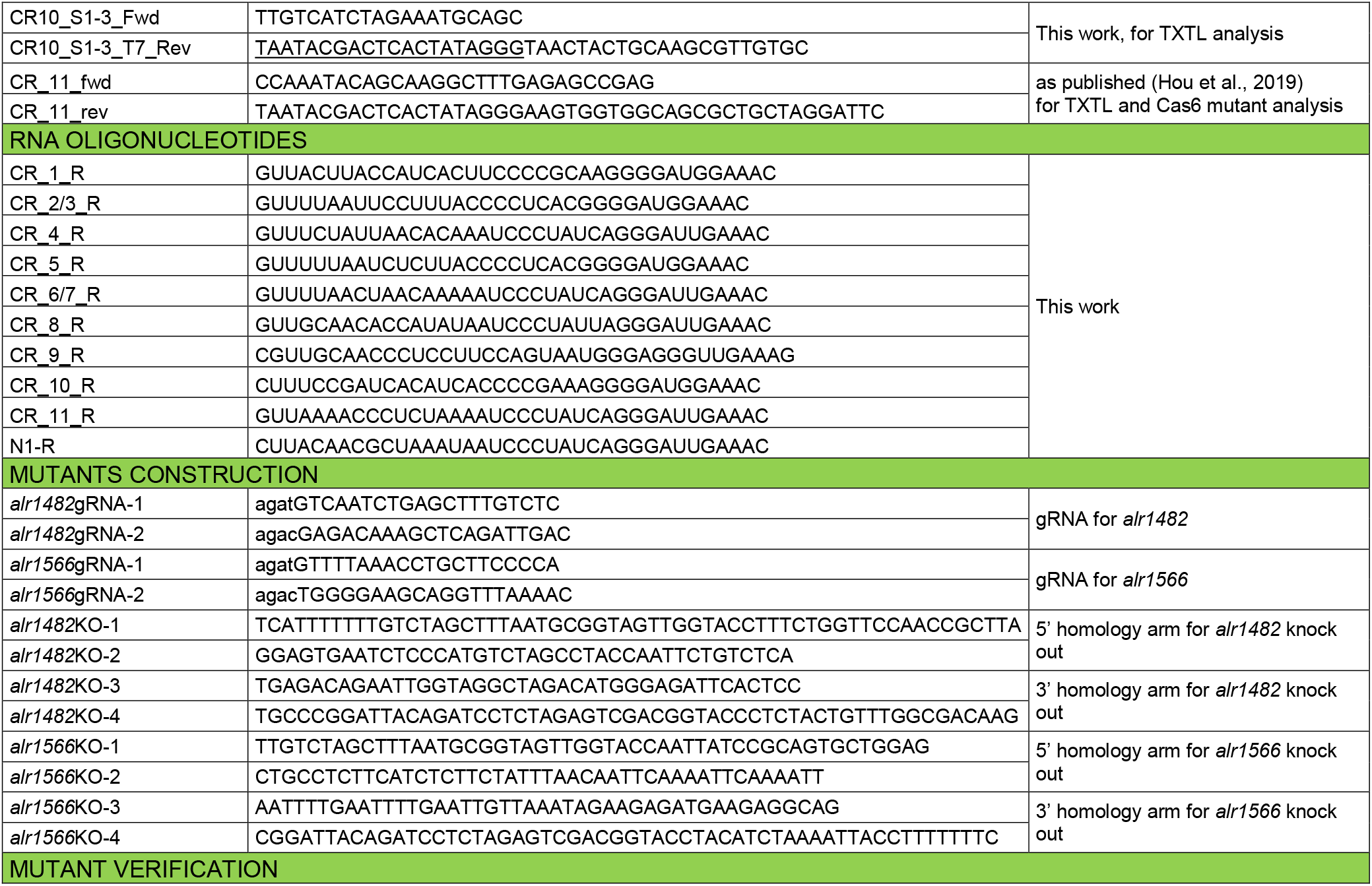
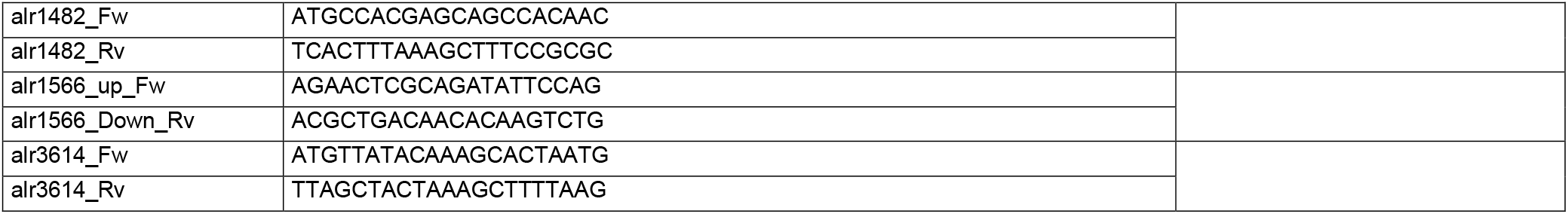
Oligonucleotide primers used in this work. Sequences belonging to the T7 promoter are underlined. Primers were purchased from IDT, RNA oligonucleotides from Biomers.

### Isolation and analysis of RNA

Total RNA was isolated from 50 mL exponentially grown cultures. Cells were harvested by centrifugation or filtration on Pall Supor 800 0.8 μm membranes. Cells were transferred to screw cap tubes together with 250 μL glass beads (0.1-0.25 μm in diameter) and 1 mL PGTX (Pinto et al., 2009) and were frozen in liquid nitrogen. Cell disruption (3 cycles of 3 x 6,500 rpm for 15 s with 10 s breaks in between) was performed using the Precellys 24 Dual homogenizer (Bertin) under nitrogen cooling. The supernatant was transferred to a new tube. Samples were incubated for 10 min at 65°C in a water bath, one volume chloroform:isoamyl alcohol (24:1) was added, and samples were incubated for 10 min at room temperature with several vortexing cycles.

After centrifugation for 3 min at 6,500 rpm in a swing-out rotor, the supernatant was transferred to a fresh tube, and one volume of chloroform:isoamyl alcohol was added. This step was repeated twice. Finally, RNA was precipitated by the addition of one volume of isopropanol. Alternatively, for the analyses of *cas6* deletion mutants, we also used the Retsch MM400 system for cell disruption. Nitrogen frozen samples were thawed and mixed with one volume of glass beads (0.1-0.25 μm and 0.5 μM in diameter, mixed in equal amounts), and cell lysis was performed in three cycles for 10 min at a frequency of 10 Hz. Between each cycle, it was checked that the samples did not become warm and were cooled if necessary. The supernatant was transferred to a fresh tube and mixed with one volume of PGTX. Then, extraction was preceded as described above.

CRISPR-related transcript accumulation was analyzed by Northern hybridization using single-stranded radioactively labeled RNA probes transcribed *in vitro* from PCR-generated templates (see **Table 1** for primers), as described (Steglich et al., 2008).

### Transcriptome analysis of the small RNA fraction of Anabaena 7120

The small RNA fraction (< 200 nt) was isolated from a single sample of total RNA from cultures grown under standard conditions using the RNeasy MinElute Cleanup kit (Qiagen). The preparation of cDNA libraries and sequencing were performed by vertis AG, Freising, Germany. The small RNA fraction sample was split into two parts. One half was first treated with Antarctic Phosphatase and rephosphorylated with T4 Polynucleotide Kinase (+ PNK), and the other half was left untreated (-PNK). Then, oligonucleotide adapters were ligated to the 5’ and 3’ ends of the RNA samples. First-strand cDNA synthesis was performed using M-MLV reverse transcriptase and the 3’ adapter as the primer. The resulting cDNAs were amplified by 12 PCR cycles using a high-fidelity DNA polymerase. The cDNA was purified using the Agencourt AMPure XP kit (Beckman Coulter) and analyzed by capillary electrophoresis. The cDNA libraries were single-end sequenced with a NextSeq 500 system using 150 bp read-length yielding 294,333,852 (+ PNK) and 36,509,792 (-PNK) reads. RNA-seq data analysis was performed with the tools installed on usegalaxy.eu. The single-end reads were trimmed, adapters and reads shorter than 14 nt were filtered out using cutadapt 1.16 (Martin, 2011). Mapping was performed on the chromosome and plasmids of *Anabaena* 7120 using bowtie2 version 2.3.4.3 with the parameters for single-end reads (Langmead and Salzberg, 2012). The data have been uploaded to NCBI’s short reads archive and were assigned the accession number PRJNA624132.

### TXTL system

For testing the Cas6 enzymes Alr1566 and Alr1482 in a cell-free transcription-translation system, the *E. coli*-based TXTL assay was used (Maxwell et al., 2018; Shin and Noireaux, 2012). The myTXTL Sigma 70 Cell-Free Master Mix was purchased from Arbor Biosciences. The included p70a plasmid was used as a template for cloning of the CRISPR repeat-spacer sequences. All PCRs were performed using PCRBio HiFi polymerase (PCR Biosystems). The p70a plasmid was linearized by PCR omitting the open reading frame for the destabilized enhanced GFP (deGFP) and its 5’UTR. CRISPR sequences were PCR-amplified from genomic DNA of *Anabaena* 7120, and overlaps with p70a were added. The repeat-spacer sequences were thus fused directly downstream of the bacteriophage λ p70a promoter and upstream of the T500 terminator.

*E. coli* K12 codon-optimized sequences of *alr1566* and *alr1482* were purchased from IDT as gBlocks and subsequently PCR-amplified with overlaps to a linearized pet28a(+) plasmid. The Cas6 proteins were thus expressed from an IPTG-inducible T7 promoter with a C-terminal 6xHis tag. Fragment assembly was performed at room temperature for 30 min upon transformation into chemically competent *E. coli* DH5alpha cells for cloning. Assembled plasmids were isolated, and regions of interest were sequenced (Eurofins Genomics).

For the expression of proteins encoded on pet28a(+) in the TXTL assay, T7 RNA polymerase, expressed in this instance from p70a, and IPTG are necessary. Reactions were performed for 16 h at 29°C in a total volume of 12 μL with 9 μL TXTL master mix, 1 mM IPTG, 0.5 nM p70a_T7RNAP, 2 nM *pet28a(+)_alr1482/alr1566*, and 10 nM p70a_CR_1-CR_11_S1-S3. For Western blot analysis, 2 μL was loaded on a 10% polyacrylamide SDS gel. Western blot analysis was performed using the Penta His HRP conjugate kit (Qiagen). To check for equal loading, the membrane was stained with Ponceau S (0.1% (w/v) in 5% acetic acid). For RNA analysis, 90 μL H2O and 100 μL phenol:chloroform:isoamyl alcohol (25:24:1) were added to 10 μL of the reactions. Phases were separated using 5Prime Phase Lock Gel heavy 2 (Quantabio). The supernatant was purified with RNA Clean and Concentrator 25 columns (Zymo Research). Ten micrograms of each sample was loaded onto a 10% polyacrylamide gel containing 8 M urea. Northern hybridizations were performed as described (Steglich et al., 2008) (see **Table 1** for probe template PCR primers).

### Expression and purification of Alr1482 and Alr1566 and cleavage assays

*E. coli* K12 codon-optimized *alr1566* and *alr1482* were expressed from pet28a(+) upon 1 mM IPTG induction in *E. coli* BL21(DE3) in a 1 L culture volume overnight at room temperature. Cell disruption (3 cycles of 3 x 6,000 rpm for 10 s interrupted by 5 s breaks) was performed with the Precellys 24 Dual homogenizer (Bertin) under nitrogen cooling. Filtered lysate was used for protein purification with an Äkta start system (GE Healthcare). Lysis buffer containing 20 mM imidazole and wash buffer with 40 mM imidazole was used. Elution was performed gradually from 40 to 500 mM imidazole. Fractions containing the purified protein were pooled and desalted with PD midi Trap G-25 (GE Healthcare) using the spin protocol, and buffer was thereby exchanged with PBS (137 mM NaCl, 2.7 mM KCl, 10 mM Na_2_HPO_4_ 1.8 mM KH_2_PO_4_; pH 7.4). Protein concentrations were determined with a direct detection spectrometer (Merck), and purified proteins were stored at −80°C. Enzymatic reactions were performed in a total reaction volume of 10 μL at 37°C for 1 h in a reaction buffer with 20 mM Tris-HCl, pH 7.8 and 400 mM KCl. As a substrate, 1 μM synthetic RNA oligonucleotides were used. The proteins Alr1482 or Alr1566 were added to a concentration of 3 μM. For experiments with ^32^P isotope-labeled probes, 30 pmol RNA was 5’-end-labeled using 40 μCi γ-ATP (3000 Ci/mmol, 10 mCi/mL, Hartmann Analytic) and 30 U of T4 polynucleotide kinase (Thermo Fisher) for 30 min at 37 °C in a volume of 30 μL. Reactions were stopped by the addition of 1.5 μL 0.5 M EDTA (pH 8.0) and incubation for 10 min at 80 °C. Unincorporated nucleotides were removed by using MicroSpin G-25 Columns (GE Healthcare). For the enzymatic reactions, the labeled RNA was incubated with 0.3 μM Cas6 at a final concentration of 0.1 μM. A total of 150 ng of the ZR small RNA ladder (Zymo Research) and 120 pmol of a homemade mix of synthetic RNA (MX with 31,29, 26, P-22 nt) were labeled analogously. To stop the reaction, RNA loading dye was added, and the fragments were separated on an 8 M urea 10% polyacrylamide sequencing gel upon staining with SYBR gold (Reimann et al., 2017). Radioactive signals were detected after drying the gel and exposing it to a storage phosphor screen (Kodak) with a GE Typhoon FLA 9500 imaging system. Synthetic CRISPR repeat oligonucleotides (**Table 1**) were purchased from Biomers.

## Results

### Two Cas6 proteins facing eight major types of repeats

Multiple sequence alignments of the ten CRISPR DRs (with the exclusion of the Tn7-associated CR_9 array) showed considerable differences between them (**Figure 1A**). As in many CRISPR-Cas systems, minor sequence variants occur (**Figure S2**). For instance, three variants exist for the CR_2 and CR_3 arrays, which belong to the same subtype III-D system but are split due to the insertion of a gene encoding Cas1-reverse transcriptase fusion protein (Hou et al., 2019). Some DRs, such as DR1 and DR7, have a sequence identity of <50% with each other (**Figure 1B**).

**Figure 1.**
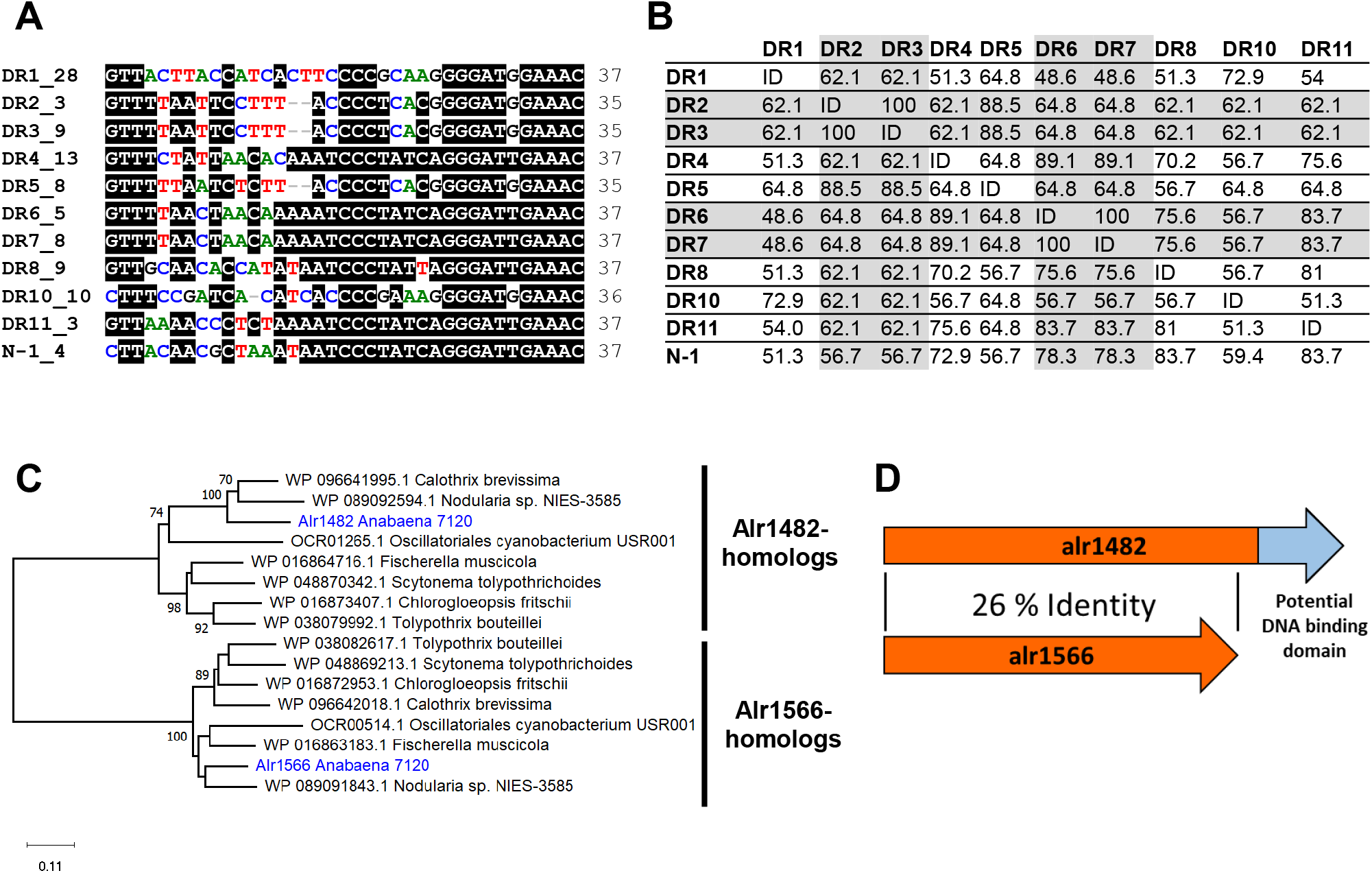
CRISPR repeats and Cas6 proteins in *Anabaena* 7120. **A.** Multiple sequence alignments of the major variants of CRISPR DRs in *Anabaena* 7120 and of cyanophage N-1, except for DR9 belonging to the Tn7-associated system. The DRs are numbered according to the respective genomic locus as defined previously (Hou et al., 2019). The numbers after the identifier give the copy number of the respective DR (e.g., CR_1 includes 28 copies of the DR major sequence variant, DR1_28). The entire alignment including all minor variants and the CR_9 Tn7-associated system is shown in **Figure S2**. **B.** Matrix comparing the DR identities against each other in %. The identical sequences of DR2 and DR3 as well as of DR6 and DR7 are shaded. **C.** Phylogenetic analysis of the two Cas6 proteins from *Anabaena* 7120 together with homologous proteins that co-occur in other selected species. Evolutionary analyses were conducted in MEGA X (Kumar et al., 2018). The complete phylogenetic analysis is shown in **Figure S3**. The phylogeny was inferred using the Minimum Evolution method (Rzhetsky and Nei, 1992). The tree with the sum of branch length = 3.299942 is shown. The percentage of replicate trees in which the associated taxa clustered together in the bootstrap test (1000 replicates) is shown next to the branches if ≥70% (Felsenstein, 1985). The tree is drawn to scale, with branch lengths in the same units as those of the evolutionary distances used to infer the phylogenetic tree. The evolutionary distances were computed using the Poisson correction method (Zuckerkandl and Pauling, 1965) and are in units of the number of amino acid substitutions per site. The ME tree was searched using the close-neighbor-interchange (CNI) algorithm (Nei and Kumar, 2000) at a search level of 1. The neighbor-joining algorithm (Saitou and Nei, 1987) was used to generate the initial tree. All ambiguous positions were removed for each sequence pair (pairwise deletion option). There were a total of 428 positions in the final dataset. **D.** Schematic depiction of both Cas6 variants of *Anabaena* 7120 in comparison.

On the other hand, some DRs are identical, such as the major variants of CR_6 and CR_7, which fits the observation that CR_6 and CR_7 belong to one and the same array and probably only recently became separated by the insertion of a 134 nt miniature inverted repeat element (MITE) (Hou et al., 2019). Hence, some of the array DRs were joined (all variants of CR_2 and CR_3 as well as CR_6 and CR_7), leaving 8 variants of clearly distinct sequences (plus the even more divergent CR_9). In addition, we included the DR of the array in phage N-1, which differed in six positions from the most closely related host-located sequence, DR8 (**Figure 1A**).

These differences raised the question of whether all these different types of DRs were substrates for one of the two Cas6 proteins, Alr1482 and Alr1566, encoded in *Anabaena* 7120, or both. These two Cas6 proteins exhibit a sequence identity of only 26% with each other and differ considerably in length from either 290 aa (Alr1566) or 377 aa (Alr1482). The extension of Alr1482 relative to Alr1566 includes an additional domain of 70 amino acids. According to the structural prediction by HHpred (Zimmermann et al., 2018), this domain possesses similarity to domain 4 of σ-factor 70, which is involved in binding to dsDNA at the −35 promoter region, as well as to domains in some repressor proteins. The possible functionality of this motif is currently elusive, but both of these similarities suggest a double-strand DNA-binding role.

Interestingly, there are close homologs to both enzymes in a large number of other cyanobacteria, and in many instances, homologs of the two proteins coexist, similar to the situation in *Anabaena* 7120. Phylogenetic analysis showed that the two Cas6 variants belong to a clearly distinct clade and a partition of all found homologs to one or the other clade (**Figure 1C** and **S3**). We identified 38 species of Cyanobacteria with at least one homolog of both types of *cas6* genes (**Figure S3**). All of these species belong to the order Nostocales, except for *Oscillatoriales cyanobacterium* USR001. Despite the general relatedness of these homologs, they varied up to 44% compared to their respective Cas6 variants in *Anabaena* 7120. The extra C-terminal domain was consistently present in all homologs in the Alr1482 clade and showed high sequence conservation.

### Analyses in a cell-free transcription-translation system

The two *Anabaena* 7120 Cas6 proteins Alr1482 and Alr1566 were expressed with a C-terminal 6xHis tag in a cell-free *E. coli*-based transcription-translation system (TXTL (Maxwell et al., 2018; Shin and Noireaux, 2012)). The expression from vector pet28a(+) was driven by T7 RNA polymerase and IPTG inducible. Both proteins were detected by Western blot analysis with respective sizes of 43 kDa for Alr1482 and 33.6 kDa for Alr1566 (**Figure 2A**). In parallel, crRNA precursors matching the sequences of CR_1, 2, 4, 5, 7, 8, 9, 10 and 11 were constitutively expressed from vector p70a. Each of these precursors consisted of the first three spacers and two DR instances (S1-R-S2-R-S3). The full-length transcripts of these different pre-crRNAs were detected by Northern hybridization (**Figure 2B**). The length of all transcripts appeared slightly higher compared to the calculated lengths, although transcription should have been terminated by the p70a terminator T500. In addition to the full-length transcripts, all reactions showed additional bands, which can often be seen for *in vitro* transcripts and is due to premature transcription termination. Nevertheless, one can clearly see that the addition of one or the other Cas6 protein changed the pattern of the pre-crRNA to shorter products with less than 100 nt. Alr1566 was able to process pre-crRNA derived from CR_4, 7, 8 and 11. Alr1482 was able to process not only CR_1,2, 5 and 10 precursors but also CR_4, 7, 8 and 11, albeit less effectively than Alr1566. The CR_9 precursor was processed by neither Alr1482 nor Alr1566. The results of these analyses are summarized in **Table 2**.

**Table 2.**
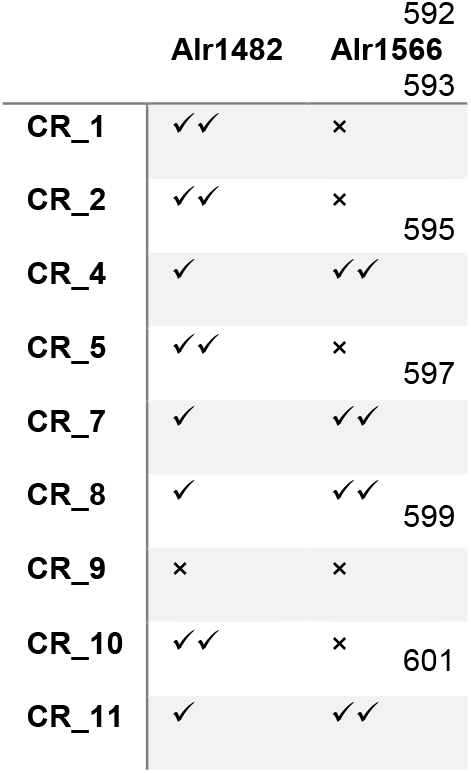
Activities of the two Cas6 enzymes in the TXTL assay (cf. Figure 2).

**Figure 2.**
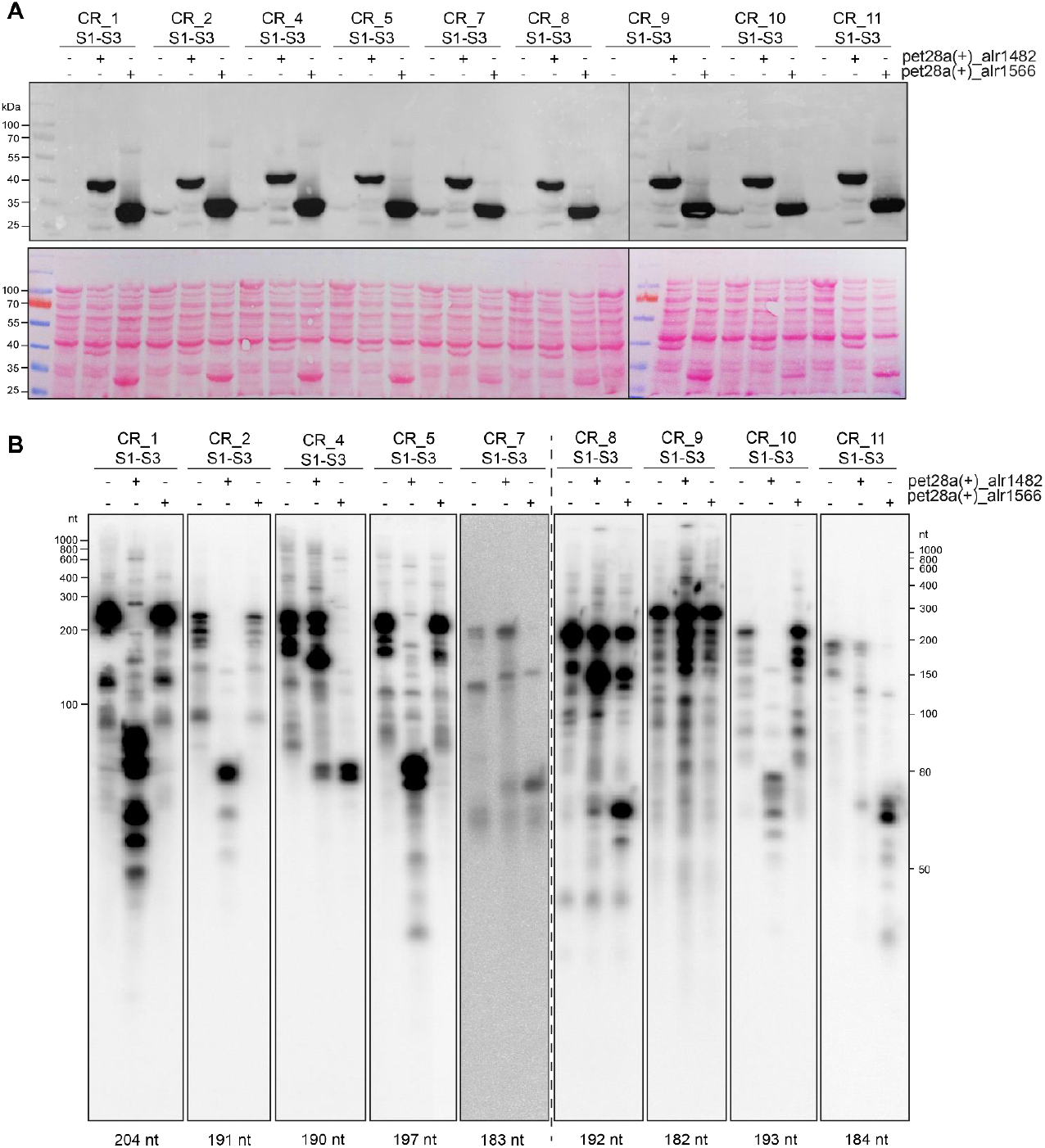
TXTL assay of Cas6 specificities. *Anabaena* 7120 Cas6 enzymes and crRNA precursors were coexpressed in a cell-free transcription-translation system (TXTL, Maxwell et al. (2018). The TXTL reactions were performed at 29°C for 16 h. **A.** Alr1566 and Alr1482 were expressed from pet28a(+). The proteins were detected by Western blot analysis via their C-terminal 6xHis tags (upper panel), and the stained membrane is shown in the lower panel. The corresponding sizes are 43 kDa for Alr1482 and 33.6 kDa for Alr1566. The prestained PageRuler (Thermo Scientific) was used as a size marker. **B.** Possible substrate transcripts encompassing the region from spacer1 to spacer3 of the respective pre-crRNAs CR_1-11 were constitutively expressed from p70a. The crRNA precursors and cleavage products were detected by Northern hybridizations using specific single-stranded RNA probes. The hypothetical length of the full-length transcript is noted underneath each hybridization panel. The size standards were Low Range ssRNA Ladder (NEB) and RiboRuler Low Range RNA Ladder (Thermo Fisher).

### Cleavage assays of CR_1-11 synthetic RNA repeats with recombinant Alr1482 or Alr1566

Both Cas6 proteins were purified as recombinant proteins and tested against all nine major DR types. **Figure 3A** shows that the DRs of CR_4, 6/7, 8 and 11 were mainly cleaved by Alr1566 but to a lesser extent by Alr1482. In contrast, the DRs of CR_1, 2/3, 5 and 10 were only cleaved by Alr1482. The DR of CR_9 was not cleaved by either of the two Cas6 proteins. For CR_1, 4, 5, 6/7 and 11, two cleavage products were observed irrespective of which of the two enzymes was responsible for the cleavage. To address the origin of the cleavage products, we performed cleavage reactions with 5’ radioactively labeled RNA, which is shown as an example for DR_1 and DR_4 in **Figure 3B**. The shorter of the two cleavage products for both repeats that were visible with SYBR Gold staining was not labeled radioactively. This finding shows that the second cleavage site was not located near the other cleavage site at the 3’ repeat end but was located close, between 2 and 4 nucleotides, to the 5’ end of the repeat.

**Figure 3.**
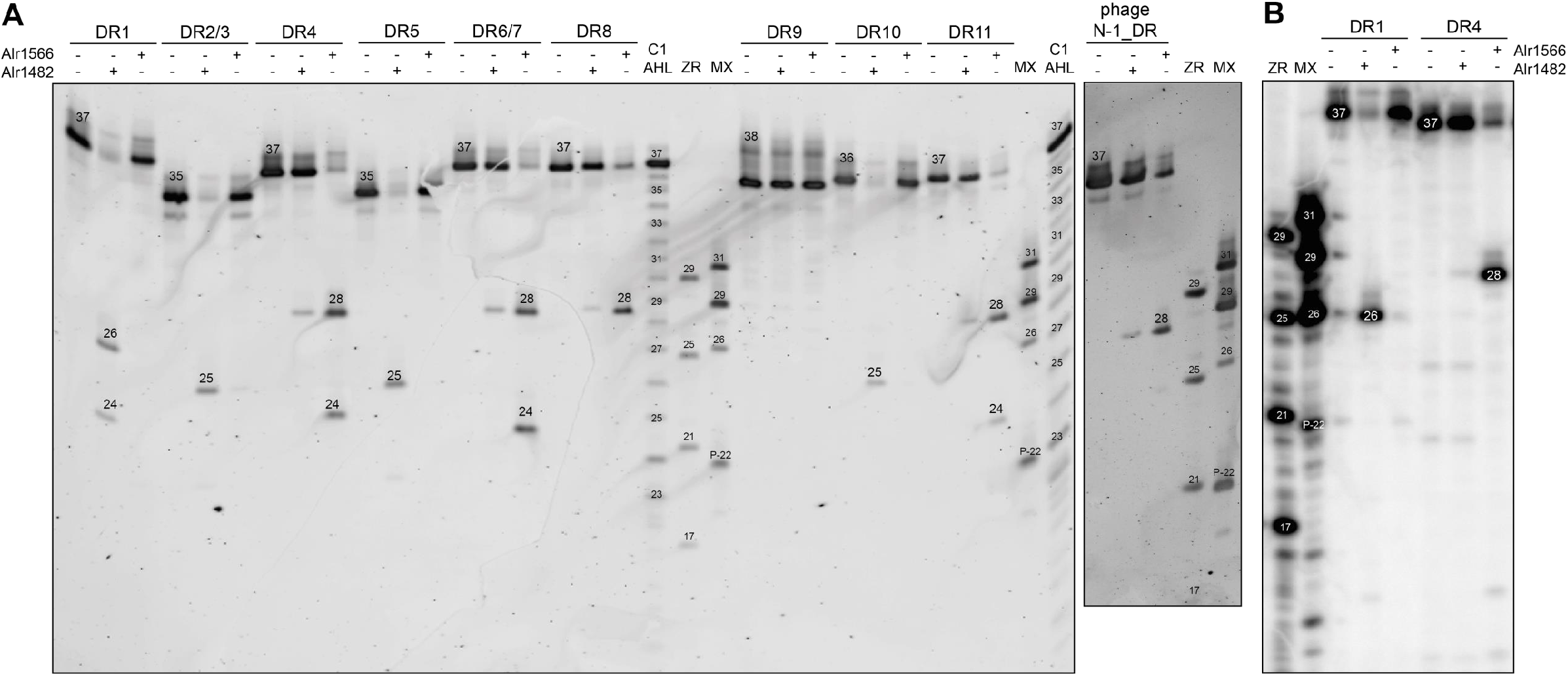
Assays using purified Cas6 proteins. **A.** Synthetic DR RNA oligonucleotides (1 μM) were incubated with the purified Cas6 proteins Alr1482 or Alr1566 (3 μM) to test for cleavage. For size estimation, the ZR small RNA ladder (Zymo Research), C1 AHL (Reimann et al., 2017) and a homemade mix of synthetic RNA (MX with 31,29, 26, P-22 nt) were used. **B**. Radioactive 5’-labeled RNA (0.1 μM) was incubated with 0.3 μM Cas6 protein. The size markers, ZR small RNA ladder (Zymo Research) and the homemade mix of synthetic RNA (MX with 31,29, 26, P-22 nt) were also 5’ radioactively labeled with γ-ATP. All samples were incubated at 37°C for 60 min, and fragments were separated on an 8 M urea 10% polyacrylamide sequencing gel. The gel shown in panel (**A**) was stained with SYBR gold, and the gel shown in panel (**B**) was dried and exposed to a storage phosphor screen. The tested RNA oligonucleotides correspond to the major DR variants, as shown in **Figure 1A**. The experiment was repeated independently three times.

The genome of cyanophage N-1 contains a CRISPR repeat-spacer array (Chénard et al., 2016). The corresponding DR was included in the *in vitro* assay with the Cas6 proteins of *Anabaena* 7120. The N-1 DR was preferentially cleaved by Alr1566 and, to a lesser extent, by Alr1482 (**Figure 3A**). Thus, the phage repeat behaved similar to the repeats of CR_4, 6/7, 8 and 11 of *Anabaena* 7120. This finding is consistent with the fact that these DR sequences shared 72.9 to 83.7% sequence identity with the N-1 DR, while all other DRs showed <60% identity (**Figure 1B**).

According to the *in vitro* cleavage results, the DRs could be classified into three groups that precisely matched the classification based on a sequence-structure comparison (**Figure 4A**). Analyses using the LocARNA algorithm that aligns sequences based on sequence and structure comparison (Raden et al., 2018; Will et al., 2012) grouped the DRs from CR_1, 2/3, 5 and 10 into one group (group 1, G1). All these DRs were cleaved by Alr1482 and not by Alr1566. The DRs of CR_4, 6/7, 8, 11 and of the N-1 phage also grouped together (group 2, G2). The DRs of G2 have in common that they were cleaved by both enzymes but more preferentially by Alr1566. In contrast to all other DRs, those of CR_9 could not be assigned to either G1 or G2 and therefore belonged to separate group 3 (G3). This classification is consistent with the TXTL results, as well. The secondary structure prediction shows a single stem-loop formed by the palindromic sequence within the DRs. The stem consists of all DRs of six to eight paired bases. An apparent difference is the nonmatching C:A in the middle of the predicted stem in all group 1 DRs (**Figure 4B**). The longer single–stranded loop region of 13 nt in DR_9 sets the only group 3 secondary structure apart from all other DRs that have loops of only three or four nucleotides (**Figure 4B**).

**Figure 4.**
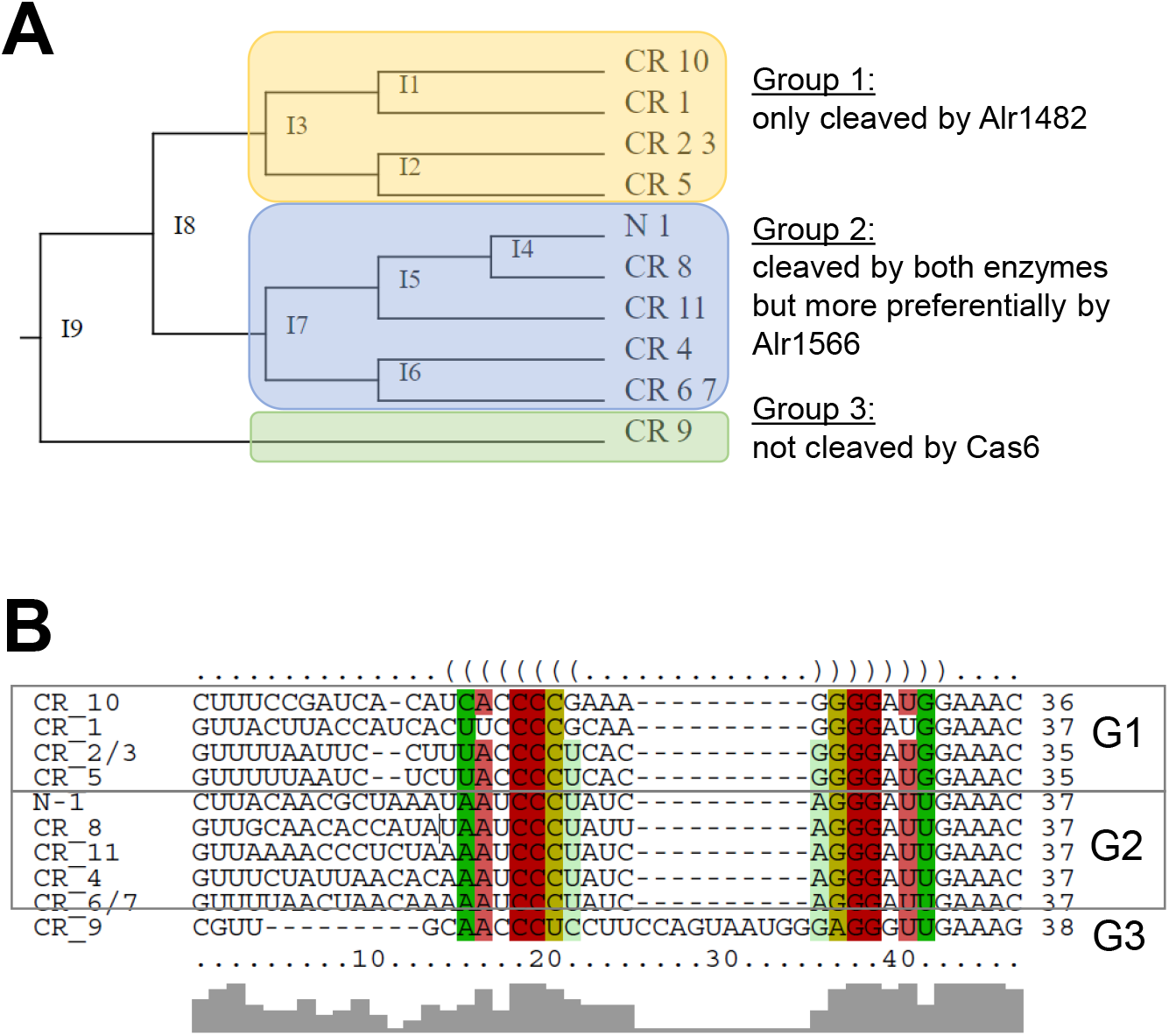
Classification of crRNA maturation in *Anabaena* 7120 according to sequence-structure alignments and Cas6 cleavage specificities. **A.** The *Anabaena* 7120 CRISPR repeats were classified by LocARNA (Raden et al., 2018; Will et al., 2012) into three different groups according to structure-sequence alignments and the generated similarity trees. This classification matched the experimentally defined groups based on the results of the *in vitro* cleavage and the TXTL reactions (**Figures 2** and **3**). Group 1 (G1) consists of DR1, 2/3, 5 and 10, and the repeats were only cleaved by Alr1482. Group 2 (G2) consists of DR4, 6/7, 8, 11 and the N-1 phage DR, and those repeats were cleaved by both enzymes but prevalently by Alr1566. DR9 is the only member of group 3 (G3) and was cleaved by none of the two Cas6 proteins. **B.** Multiple structure-sequence alignment of all *Anabaena* 7120 DRs with annotation of the predicted secondary structures.

### Analysis of cas6 deletion mutants provides in vivo evidence for crosstalk

Cas6 enzymes generate the crRNA 5’ tags, in many systems by a single cleavage eight nucleotides from the 3’ end of the respective repeat (for review see Behler and Hess (2019). The results of the *in vitro* assays shown in **Figure 3** seem to deviate from this rule because in several instances, two bands were generated. Moreover, the longer cleavage fragments differed in lengths, seemingly one or two nucleotides too short to leave 8 nucleotide-long tags. Hence, to compare the results of the *in vitro* and *in silico* analyses with the *in vivo* situation, an sRNA transcriptome analysis of WT cultures was performed and deletion mutants of the two *cas6* genes (*Δalr1482* and *Δalr1566*) were constructed and analyzed. Consistent with the *in vitro* data, the crRNAs of all group 1 arrays were expressed and processed in the WT and Δ *alr1566* but not in *Δalr1482*, as shown in **Figure 5A**, for this group using the CR_1 array as an example. Moreover, the data from the sRNA transcriptome analysis of WT cultures showed that *in vivo*, the mature crRNAs started precisely with the 8 nt 5’ tags that are needed for self/nonself discrimination and loading of the mature crRNA into the effector complexes (**Figure 5B**). Similar results were obtained for the other group 1 DRs (**Figures S5 to S7**).

**Figure 5.**
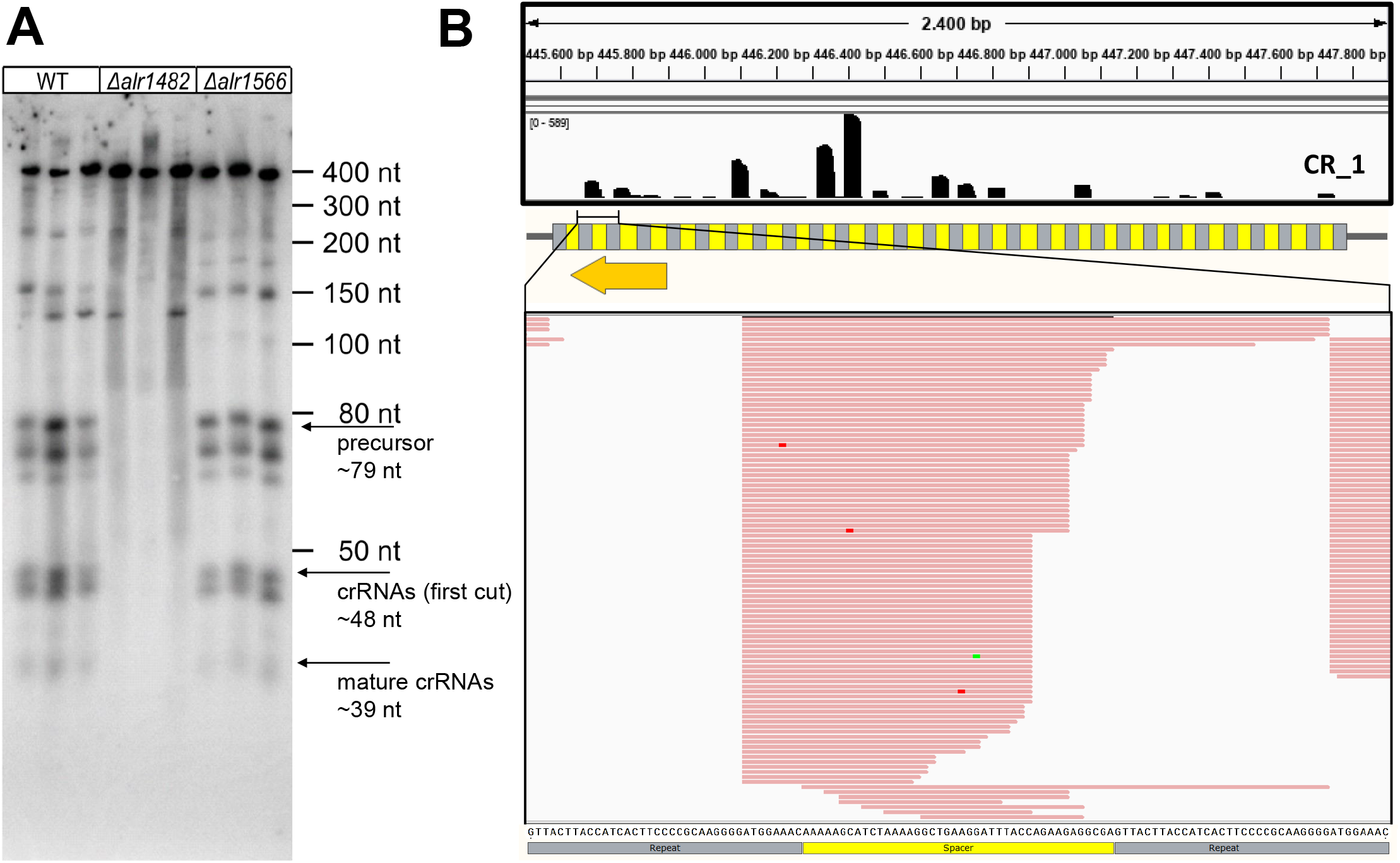
Analysis of *cas6* deletion mutants for the CR_1 array representing group 1 arrays *in vivo*. **A.** The accumulation of crRNAs and precursors for the CR_1 repeat-spacer array after RNA separation in high-resolution polyacrylamide gels and Northern hybridization. The blots show RNA hybridization results with three clones each from WT and the deletion mutants *Δalr1482* and *Δalr1566*. **B.** On top, the coverage in a small RNA transcriptome in WT cells is shown along the full length of the respective array (gray, repeats; yellow, spacers) and below in detail for a selected number of reads. The color of the reads highlights their direction (red: forward; blue: reverse), and mutations inside the reads are marked by different colors (adenine: green; cytosine: blue; guanine: orange; thymine: red). The sequence below represents the DNA sequence of the sense strand of the CRISPR array. The region recognized by the respective probe in the Northern hybridization in panel (A) is indicated by the orange arrow. The orientation of the CRISPR array is illustrated by a black arrow.

In contrast, the CRISPR arrays of group 2 were processed just in the WT and in *Δalr1482*, while in *Δalr1566*, only some precursors >200 nt in length were detected, exemplified for this group by means of the CR_8 array in **Figure 6A**. Again, the comparison to sRNA transcriptome data from WT cultures showed that *in vivo*, the mature crRNAs started precisely with the 8 nt 5’ tags *in vivo* (**Figure 6B**).

**Figure 6.**
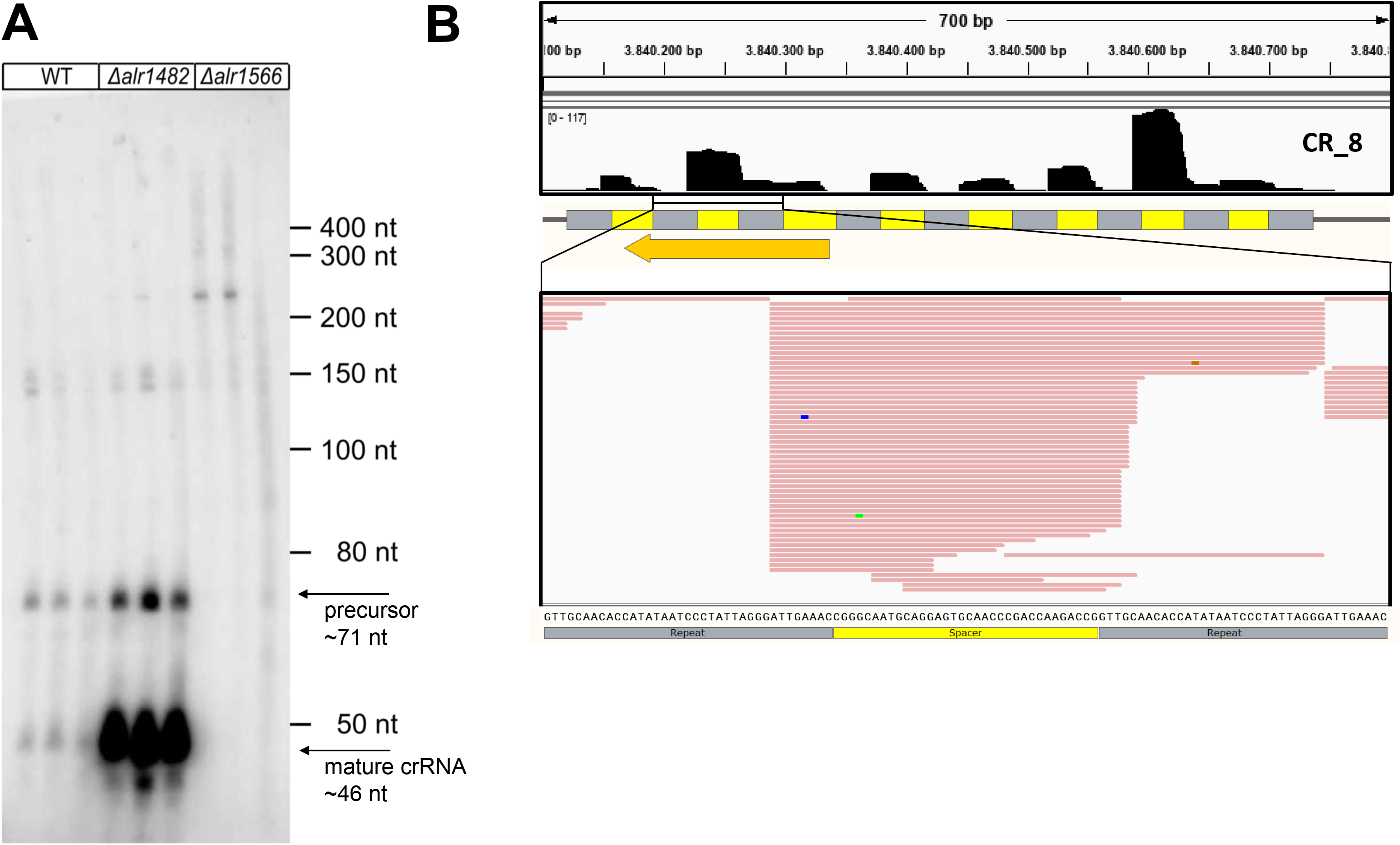
Analysis of *cas6* deletion mutants for the CR_8 array representing group 2 arrays *in vivo*. **A.** The accumulation of crRNAs and precursors for the CR_8 repeat-spacer array after RNA separation in high-resolution polyacrylamide gels and Northern hybridization. The blots show RNA hybridization results with three clones each from WT and the deletion mutants *Δalr1482* and *Δalr1566*. **B.** On top, the coverage in a small RNA transcriptome in WT cells is shown along the full length of the respective array (gray, repeats; yellow, spacers) and below in detail for a selected number of reads. The color of the reads highlights their direction (red: forward; blue: reverse), and mutations inside the reads are marked by different colors (adenine: green; cytosine: blue; guanine: orange; thymine: red). The sequence below represents the DNA sequence of the sense strand of the CRISPR array. The region recognized by the respective probe in the Northern hybridization in panel (A) is indicated by the orange arrow. The orientation of the CRISPR array is illustrated by a black arrow.

These data suggest that Alr1482 is not able to properly cleave the group 2 repeats *in vivo*. However, a striking observation was the highly increased accumulation of mature CR_8 crRNAs in *Δalr1482* compared to the WT. This overaccumulation is also visible for the other group 2 crRNAs (**Figures S8** to **S10**).

## Discussion

While the coexistence of multiple CRISPR-Cas systems in one organism is frequent, there are notably few studies in which the role of co-occurring Cas6 was investigated. Reimann et al. (2017) found evidence for weak promiscuity of Cas6-1 and Cas6-2a in the cyanobacterium *Synechocystis* sp. PCC 6803 *in vitro* but not *in vivo*. In the archaeon *Pyrococcus furiosus*, three CRISPR-Cas systems exist (Types I-A, I-G and III-B) and seven repeat-spacer arrays, while only a single *cas6* gene is present (Majumdar et al., 2015). In another Euryarchaeon, *Methanosarcina mazei* strain Gö1, a Type I-B and a Type III-C system coexist, with each being associated with its own CRISPR array and a single *cas6* gene (Nickel et al., 2013). Assays *in vitro* unexpectedly indicated that both Cas6 were able to process the precursor RNA from both CRISPR arrays, while *in vivo*, only one of the enzymes, that of the I-B system, mediated the maturation of crRNAs from both systems (Nickel et al., 2019). In contrast to observations made in *Anabaena* 7120, the DR sequences in *Methanosarcina mazei* Gö1 are highly similar and differ by a maximum of two positions.

In our specificity assays, we included the major DR sequence of the CRISPR array in cyanophage N-1 (**Figure 3A**). This sequence was cleaved by both enzymes but more preferentially by Alr1566, consistent with its assignment to group 2 according to the classification based on sequence and secondary structure (**Figure 4**). These data strongly suggest that the *cas* gene-free system of phage N-1 is fully competent to interact with host-encoded Cas proteins. While there are no reliable targets predicted for the four spacers present in the N-1 array, we identified a single spacer in *Anabaena* 7120 to target the N-1 genome (**Figure S4**). Spacer 12 of the CR_1 system is identical in 40 of its 45 nt positions to the region from nucleotides 5,214 to 5,258 on the reverse strand of the N-1 genome (GenBank accession KU234532), in which an unknown protein is encoded that in database searches revealed a single homolog in cyanophage A-1. These findings further support the tight interaction between this phage and *Anabaena* 7120. However, the five mismatches between CR_1 spacer 12 and the phage N-1 genome likely render the immunity inactive because we observed a high efficiency of plaque formation (not shown).

Our data provide further evidence that all the distributed CRISPR arrays in *Anabaena* 7120 are likely functional. The results clearly assign all 11 identified repeat spacer arrays to three systems:

- group 1 crRNAs, which result *in vivo* and *in vitro* only from processing by Alr1482, belong to the Type III-D system,
- group 2 crRNAs, which are primarily processed by Alr1566, belong to the Type I-D system
- the group 3 array belongs to the Tn7-associated system and is independent of either Cas6.

The most striking finding was the group 2 arrays that are primarily processed by Alr1566 and overaccumulated in the *Δalr1482* mutant compared to the WT (**Figure 6, Figure S8 to S10**). The fully matured crRNAs are the key element for the assembly of functionally active defense complexes. Therefore, it is likely that in this condition, more functional CRISPR-Cas complexes exist that carry group 2 crRNAs. This property probably is biologically meaningful, for example, when the other type of complexes, in this instance, the Type III-D system carrying group 1 crRNAs, would be inactivated. Such inactivation could result from mutations or neutralization by an anti-CRISPR factor. Anti-CRISPR proteins have been characterized for several different types of CRISPR-Cas systems and have also been predicted for Type III-D systems (Gussow et al., 2020).

The main conclusion from these observations is that the parallel presence of Alr1482 in the WT normally has a dampening effect on the expression of the Type I-D system carrying group 2 crRNAs. This effect could be mediated at the posttranscriptional level. In this scenario, the observed weak cross-reactivity of Alr1482 for group 2 crRNA maturation (**Figure 3A**) might lead to an association of the Alr1482 Cas6 endonuclease with the respective products of this maturation. The resulting group 2 crRNA-Alr1482 complexes are likely incompatible with the assembly of the I-D interference complexes and therefore become rapidly degraded. In an alternative scenario, this crosstalk could be mediated directly via the potential DNA binding domain in Alr1482 (**Figure 1D**). Clarifying the functional relevance of this domain as well as identifying the evolutionary forces and genetic mechanisms driving the frequent fragmentation of CRISPR-Cas repeat-spacer arrays in filamentous cyanobacteria, warrant further research.

## Funding information

Financial support for this work was provided by the Joint Sino-German Research Program (grant HE 2544/13-1 to WRH), the National Natural Science Foundation of China (grant 31761133008 to XL), the QIBEBT and Dalian National Laboratory for Clean Energy (DNL), CAS (grant QIBEBT I201904 to TZ), and the Shandong Taishan Scholarship (to XL).

## Author contributions

VR, TZ and MZ performed the experimental analyses, and bioinformatics analyses were performed by MZ and WRH. HL contributed to the strain construction, and JB contributed to the small RNA analysis. WRH, VR, TZ and XL drafted the manuscript with contributions from all authors.

## Notes

The authors have no conflicts of interest to declare.

### Competing Interest Statement

The authors have declared no competing interest.

## References

Adolph, K.W., and Haselkorn, R. (1971). Isolation and characterization of a virus infecting the blue-green alga *Nostoc muscorum*. Virology 46, 200–208.

Adolph, K.W., and Haselkorn, R. (1973). Blue-green algal virus N-1: Physical properties and disassembly into structural parts. Virology 53, 427–440.

Behler, J., and Hess, W.R. (2019). Approaches to study CRISPR RNA biogenesis and the key players involved. Methods 172, 12–26.

Behler, J., Sharma, K., Reimann, V., Wilde, A., Urlaub, H., and Hess, W.R. (2018). The host-encoded RNase E endonuclease as the crRNA maturation enzyme in a CRISPR–Cas subtype III-Bv system. Nature Microbiology 3, 367–377.

Carte, J., Wang, R., Li, H., Terns, R.M., and Terns, M.P. (2008). Cas6 is an endoribonuclease that generates guide RNAs for invader defense in prokaryotes. Genes Dev 22, 3489–3496.

Charpentier, E., Richter, H., van der Oost, J., and White, M.F. (2015). Biogenesis pathways of RNA guides in archaeal and bacterial CRISPR-Cas adaptive immunity. FEMS Microbiol Rev 39, 428–441.

Chénard, C., Wirth, J.F., and Suttle, C.A. (2016). Viruses infecting a freshwater filamentous Cyanobacterium (*Nostoc* sp.) encode a functional CRISPR array and a proteobacterial DNA polymerase B. MBio 7, e00667–16.

Chou-Zheng, L., and Hatoum-Aslan, A. (2019). A type III-A CRISPR-Cas system employs degradosome nucleases to ensure robust immunity. Elife 8, e45393.

Elhai, J., and Wolk, C.P. (1988). Conjugal transfer of DNA to cyanobacteria. Meth Enzymol 167, 747–754.

Felsenstein, J. (1985). Confidence limits on phylogenies: An approach using the bootstrap. Evolution 39, 783–791.

Gussow, A.B., Shmakov, S.A., Makarova, K.S., Wolf, Y.I., Bondy-Denomy, J., and Koonin, E.V. (2020). Vast diversity of anti-CRISPR proteins predicted with a machine-learning approach. BioRxiv 2020.01.23.916767.

Hale, C., Kleppe, K., Terns, R.M., and Terns, M.P. (2008). Prokaryotic silencing (psi)RNAs in *Pyrococcus furiosus*. RNA 14, 2572–2579.

Hale, C.R., Zhao, P., Olson, S., Duff, M.O., Graveley, B.R., Wells, L., Terns, R.M., and Terns, M.P. (2009). RNA-guided RNA cleavage by a CRISPR RNA-Cas protein complex. Cell 139, 945–956.

Hochstrasser, M.L., and Doudna, J.A. (2015). Cutting it close: CRISPR-associated endoribonuclease structure and function. Trends Biochem. Sci. 40, 58–66.

Hou, S., Brenes-Álvarez, M., Reimann, V., Alkhnbashi, O.S., Backofen, R., Muro-Pastor, A.M., and Hess, W.R. (2019). CRISPR-Cas systems in multicellular cyanobacteria. RNA Biol 16, 518–529.

Karginov, F.V., and Hannon, G.J. (2010). The CRISPR system: small RNA-guided defense in Bacteria and Archaea. Molecular Cell 37, 7–19.

Kumar, S., Stecher, G., Li, M., Knyaz, C., and Tamura, K. (2018). MEGA X: Molecular evolutionary genetics analysis across computing platforms. Mol Biol Evol 35, 1547–1549.

Langmead, B., and Salzberg, S.L. (2012). Fast gapped-read alignment with Bowtie 2. Nat Meth 9, 357–359.

Madeira, F., Park, Y. mi, Lee, J., Buso, N., Gur, T., Madhusoodanan, N., Basutkar, P., Tivey, A.R.N., Potter, S.C., Finn, R.D., et al. (2019). The EMBL-EBI search and sequence analysis tools APIs in 2019. Nucleic Acids Res 47, W636–W641.

Majumdar, S., Zhao, P., Pfister, N.T., Compton, M., Olson, S., Glover, C.V.C., Wells, L., Graveley, B.R., Terns, R.M., and Terns, M.P. (2015). Three CRISPR-Cas immune effector complexes coexist in *Pyrococcus furiosus*. RNA 21, 1147–1158.

Makarova, K.S., Wolf, Y.I., Alkhnbashi, O.S., Costa, F., Shah, S.A., Saunders, S.J., Barrangou, R., Brouns, S.J.J., Charpentier, E., Haft, D.H., et al. (2015). An updated evolutionary classification of CRISPR-Cas systems. Nat Rev Microbiol 13, 722–736.

Martin, M. (2011). Cutadapt removes adapter sequences from high-throughput sequencing reads. EMBnet. Journal 17, 10.

Maxwell, C.S., Jacobsen, T., Marshall, R., Noireaux, V., and Beisel, C.L. (2018). A detailed cell-free transcription-translation-based assay to decipher CRISPR protospacer-adjacent motifs. Methods 143, 48–57.

Nei, M., and Kumar, S. (2000). Molecular Evolution and Phylogenetics (New York, NY: Oxford University Press).

Nickel, L., Weidenbach, K., Jäger, D., Backofen, R., Lange, S.J., Heidrich, N., and Schmitz, R.A. (2013). Two CRISPR-Cas systems in *Methanosarcina mazei* strain Gö1 display common processing features despite belonging to different types I and III. RNA Biol 10, 779–791.

Nickel, L., Ulbricht, A., Alkhnbashi, O.S., Förstner, K.U., Cassidy, L., Weidenbach, K., Backofen, R., and Schmitz, R.A. (2019). Cross-cleavage activity of Cas6b in crRNA processing of two different CRISPR-Cas systems in *Methanosarcina mazei* Gö1. RNA Biol 16, 492–503.

Pinto, F.L., Thapper, A., Sontheim, W., and Lindblad, P. (2009). Analysis of current and alternative phenol based RNA extraction methodologies for cyanobacteria. BMC Molecular Biology 10, 79.

Przybilski, R., Richter, C., Gristwood, T., Clulow, J.S., Vercoe, R.B., and Fineran, P.C. (2011). Csy4 is responsible for CRISPR RNA processing in *Pectobacterium atrosepticum*. RNA Biology 8, 517–528.

Raden, M., Ali, S.M., Alkhnbashi, O.S., Busch, A., Costa, F., Davis, J.A., Eggenhofer, F., Gelhausen, R., Georg, J., Heyne, S., et al. (2018). Freiburg RNA tools: a central online resource for RNA-focused research and teaching. Nucleic Acids Res. 46, W25–W29.

Reimann, V., Alkhnbashi, O.S., Saunders, S.J., Scholz, I., Hein, S., Backofen, R., and Hess, W.R. (2017). Structural constraints and enzymatic promiscuity in the Cas6-dependent generation of crRNAs. Nucleic Acids Research 45, 915–925.

Rippka, R., Deruelles, J., Waterbury, J.B., Herdman, M., and Stanier, R.Y. (1979). Generic assignments, strain histories and properties of pure cultures of cyanobacteria. Microbiology 111, 1–61.

Rzhetsky, A., and Nei, M. (1992). A simple method for estimating and testing minimum evolution trees. Mol Biol Evol 9, 945–967.

Saitou, N., and Nei, M. (1987). The neighbor-joining method: A new method for reconstructing phylogenetic trees. Mol Biol Evol 4, 406–425.

Shah, S.A., Alkhnbashi, O.S., Behler, J., Han, W., She, Q., Hess, W.R., Garrett, R.A., and Backofen, R. (2019). Comprehensive search for accessory proteins encoded with archaeal and bacterial type III CRISPR-cas gene cassettes reveals 39 new cas gene families. RNA Biol 16, 530–542.

Shin, J., and Noireaux, V. (2012). An *E. coli* cell-free expression toolbox: application to synthetic gene circuits and artificial cells. ACS Synth Biol 1, 29–41.

Steglich, C., Futschik, M.E., Lindell, D., Voss, B., Chisholm, S.W., and Hess, W.R. (2008). The challenge of regulation in a minimal photoautotroph: non-coding RNAs in *Prochlorococcus*. PLOS Genetics 4, e1000173.

Strecker, J., Ladha, A., Gardner, Z., Schmid-Burgk, J.L., Makarova, K.S., Koonin, E.V., and Zhang, F. (2019). RNA-guided DNA insertion with CRISPR-associated transposases. Science 365, 48–53.

Ungerer, J., and Pakrasi, H.B. (2016). Cpf1 is a versatile tool for CRISPR genome editing across diverse species of Cyanobacteria. Sci Rep 6, 39681.

Will, S., Joshi, T., Hofacker, I.L., Stadler, P.F., and Backofen, R. (2012). LocARNA-P: accurate boundary prediction and improved detection of structural RNAs. RNA 18, 900–914.

Zhang, Q., and Ye, Y. (2017). Not all predicted CRISPR–Cas systems are equal: isolated cas genes and classes of CRISPR like elements. BMC Bioinformatics 18, 92.

Zimmermann, L., Stephens, A., Nam, S.-Z., Rau, D., Kübler, J., Lozajic, M., Gabler, F., Söding, J., Lupas, A.N., and Alva, V. (2018). A completely reimplemented MPI bioinformatics toolkit with a new HHpred server at its core. J Mol Biol 430, 2237–224.

Zuckerkandl, E., and Pauling, L. (1965). Evolutionary divergence and convergence in proteins. In Evolving Genes and Proteins, V. Bryson, and H.J. Vogel, eds. (New York, NY: Academic Press), pp. 97–166.

