## Supplementary Figures S1 to S10 for "Specificities of and functional coordination between the two Cas6 maturation endonucleases in *Anabaena* sp. PCC 7120 assign orphan CRISPR arrays to three groups"

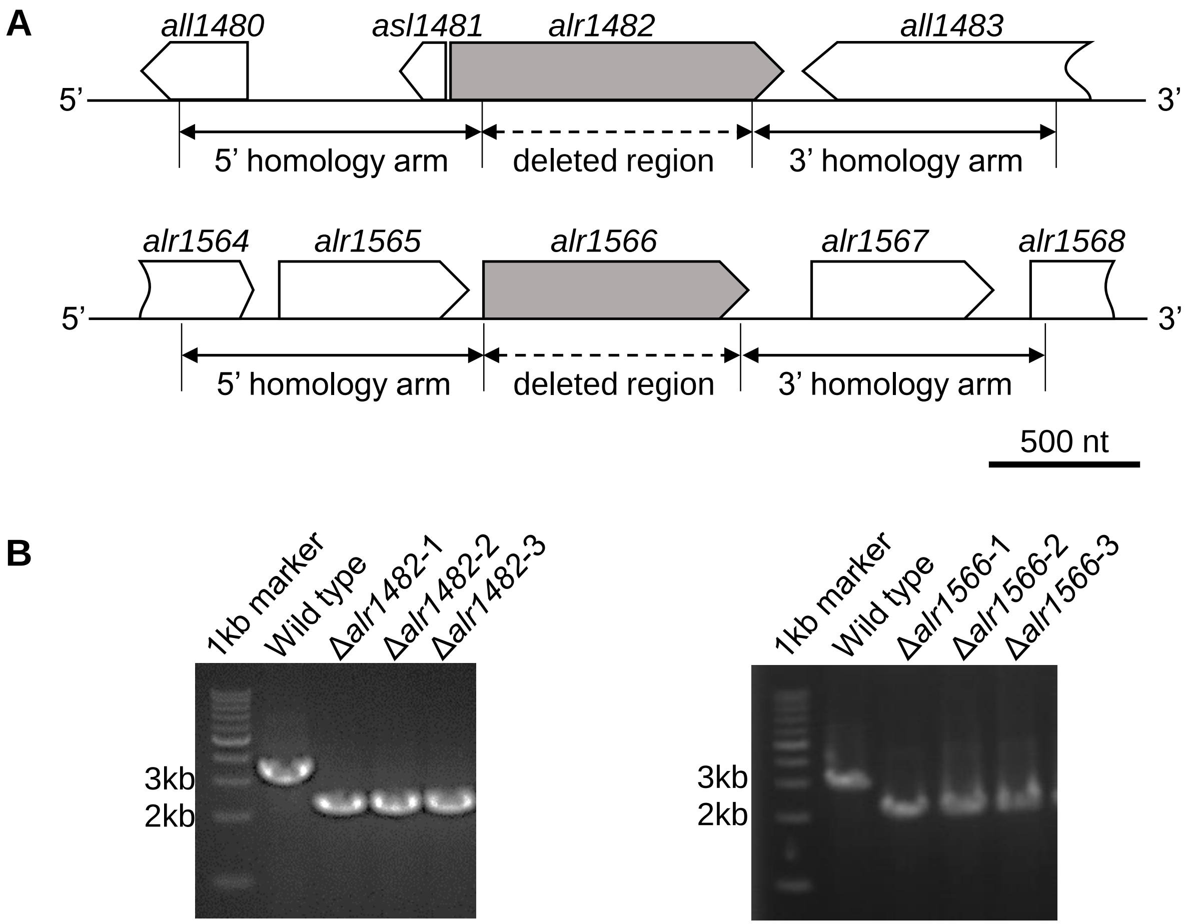


**Figure S1. Construction of *cas6* deletion mutants in *Anabaena* 7120. A.** Schematic drawing of the deleted regions and flanking regions. **B.** PCR assays showing successful segregation of deletion mutants in three independent replicates each.

**A**

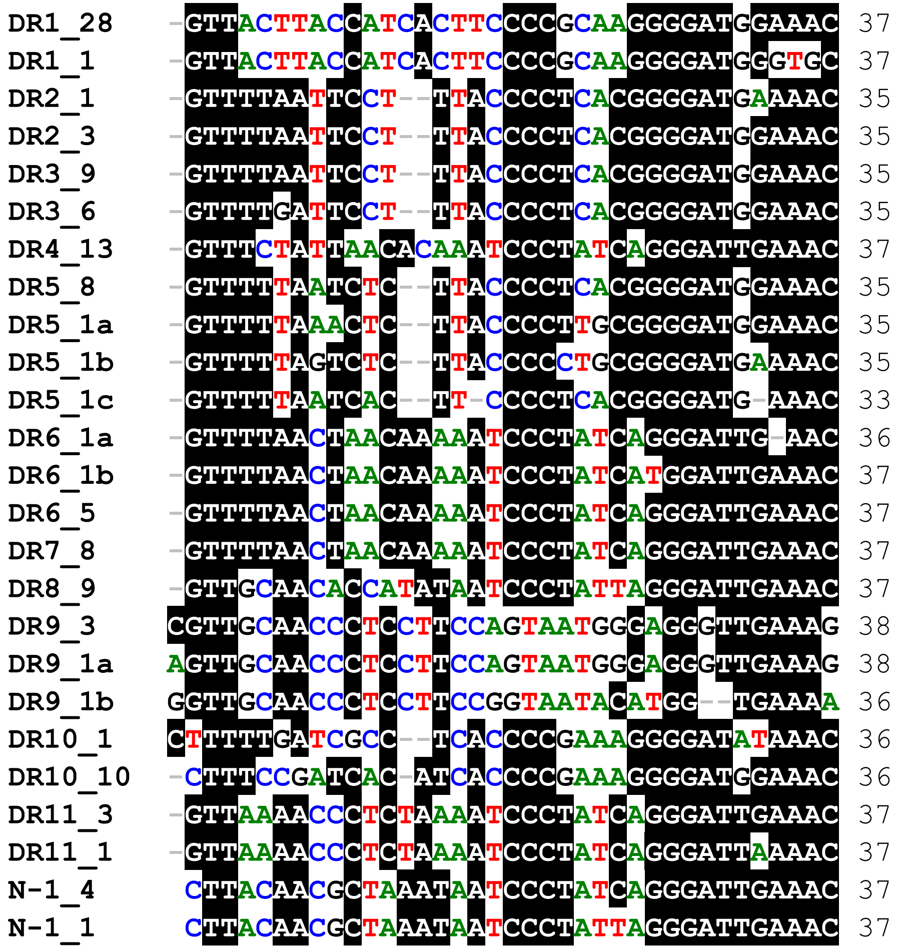


**B**

| | DR1_28 | DR1_1 | DR2_1 | DR2_3 | DR3_9 | DR3_6 | DR4_13 | DR5_8 | DR5_1a | DR5_1b | DR5_1c | DR6_1a | DR6_1b | DR6_5 | DR7_8 | DR8_9 | DR9_3 | DR9_1a | DR9_1b | DR10_1 | DR10_10 | DR11_3 | DR11_1 | N1_4 | N1_1 |
|---|---|---|---|---|---|---|---|---|---|---|---|---|---|---|---|---|---|---|---|---|---|---|---|---|---|
| **DR1_28** | ID | 918 | 621 | 648 | 648 | 648 | 513 | 675 | 621 | 567 | 675 | 459 | 459 | 486 | 486 | 513 | 394 | 394 | 368 | 526 | 675 | 540 | 513 | 513 | 513 |
| **DR1_1** | 918 | ID | 540 | 567 | 567 | 567 | 432 | 594 | 540 | 486 | 594 | 405 | 378 | 405 | 405 | 432 | 315 | 315 | 289 | 447 | 594 | 459 | 432 | 432 | 432 |
| **DR2_1** | 621 | 540 | ID | 971 | 971 | 942 | 594 | 857 | 771 | 800 | 828 | 594 | 594 | 621 | 621 | 594 | 368 | 368 | 368 | 666 | 638 | 567 | 594 | 567 | 540 |
| **DR2_3** | 648 | 567 | 971 | ID | 1000 | 971 | 621 | 885 | 800 | 771 | 828 | 621 | 621 | 648 | 648 | 621 | 394 | 394 | 394 | 666 | 666 | 594 | 567 | 594 | 567 |
| **DR3_9** | 648 | 567 | 971 | 1000 | ID | 971 | 621 | 885 | 800 | 771 | 828 | 621 | 621 | 648 | 648 | 621 | 394 | 394 | 394 | 666 | 666 | 594 | 567 | 594 | 567 |
| **DR3_6** | 648 | 567 | 942 | 971 | 971 | ID | 621 | 885 | 800 | 771 | 828 | 594 | 594 | 621 | 621 | 594 | 368 | 368 | 368 | 694 | 666 | 567 | 540 | 567 | 540 |
| **DR4_13** | 513 | 432 | 594 | 621 | 621 | 621 | ID | 648 | 621 | 594 | 621 | 864 | 864 | 891 | 891 | 702 | 368 | 368 | 394 | 526 | 621 | 756 | 729 | 729 | 702 |
| **DR5_8** | 675 | 594 | 857 | 885 | 885 | 885 | 648 | ID | 914 | 857 | 914 | 621 | 621 | 648 | 648 | 567 | 421 | 421 | 421 | 638 | 722 | 621 | 594 | 594 | 567 |
| **DR5_1a** | 621 | 540 | 771 | 800 | 800 | 800 | 621 | 914 | ID | 885 | 828 | 594 | 594 | 621 | 621 | 594 | 473 | 473 | 421 | 611 | 666 | 621 | 594 | 594 | 567 |
| **DR5_1b** | 567 | 486 | 800 | 771 | 771 | 771 | 594 | 857 | 885 | ID | 800 | 567 | 567 | 594 | 594 | 513 | 447 | 447 | 394 | 611 | 638 | 567 | 594 | 540 | 513 |
| **DR5_1c** | 675 | 594 | 828 | 828 | 828 | 828 | 621 | 914 | 828 | 800 | ID | 594 | 594 | 621 | 621 | 513 | 368 | 368 | 368 | 611 | 694 | 540 | 540 | 513 | 486 |
| **DR6_1a** | 459 | 405 | 594 | 621 | 621 | 594 | 864 | 621 | 594 | 567 | 594 | ID | 945 | 972 | 972 | 729 | 368 | 368 | 394 | 500 | 567 | 810 | 783 | 756 | 729 |
| **DR6_1b** | 459 | 378 | 594 | 621 | 621 | 594 | 864 | 621 | 594 | 567 | 594 | 945 | ID | 972 | 972 | 729 | 394 | 394 | 447 | 500 | 567 | 810 | 783 | 756 | 729 |
| **DR6_5** | 486 | 405 | 621 | 648 | 648 | 621 | 891 | 648 | 621 | 594 | 621 | 972 | 972 | ID | 1000 | 756 | 394 | 394 | 421 | 526 | 594 | 837 | 810 | 783 | 756 |
| **DR7_8** | 486 | 405 | 621 | 648 | 648 | 621 | 891 | 648 | 621 | 594 | 621 | 972 | 972 | 1000 | ID | 756 | 394 | 394 | 421 | 526 | 594 | 837 | 810 | 783 | 756 |
| **DR8_9** | 513 | 432 | 594 | 621 | 621 | 594 | 702 | 567 | 594 | 513 | 513 | 729 | 729 | 756 | 756 | ID | 473 | 473 | 473 | 500 | 540 | 810 | 783 | 837 | 864 |
| **DR9_3** | 394 | 315 | 368 | 394 | 394 | 368 | 368 | 421 | 473 | 447 | 368 | 368 | 394 | 394 | 394 | 473 | ID | 973 | 763 | 394 | 394 | 473 | 447 | 447 | 447 |
| **DR9_1a** | 394 | 315 | 368 | 394 | 394 | 368 | 368 | 421 | 473 | 447 | 368 | 368 | 394 | 394 | 394 | 473 | 973 | ID | 763 | 368 | 394 | 473 | 447 | 447 | 447 |
| **DR9_1b** | 368 | 289 | 368 | 394 | 394 | 368 | 394 | 421 | 421 | 394 | 368 | 394 | 447 | 421 | 421 | 473 | 763 | 763 | ID | 342 | 368 | 500 | 473 | 473 | 447 |
| **DR10_1** | 526 | 447 | 666 | 666 | 666 | 694 | 526 | 638 | 611 | 611 | 611 | 500 | 500 | 526 | 526 | 500 | 394 | 368 | 342 | ID | 675 | 500 | 500 | 473 | 473 |
| **DR10_10** | 675 | 594 | 638 | 666 | 666 | 666 | 621 | 722 | 666 | 638 | 694 | 567 | 567 | 594 | 594 | 540 | 394 | 394 | 368 | 675 | ID | 513 | 486 | 594 | 594 |
| **DR11_3** | 540 | 459 | 567 | 594 | 594 | 567 | 756 | 621 | 621 | 567 | 540 | 810 | 810 | 837 | 837 | 810 | 473 | 473 | 500 | 500 | 513 | ID | 972 | 837 | 810 |
| **DR11_1** | 513 | 432 | 594 | 567 | 567 | 540 | 729 | 594 | 594 | 594 | 540 | 783 | 783 | 810 | 810 | 783 | 447 | 447 | 473 | 500 | 486 | 972 | ID | 810 | 783 |
| **N1_4** | 513 | 432 | 567 | 594 | 594 | 567 | 729 | 594 | 594 | 540 | 513 | 756 | 756 | 783 | 783 | 837 | 447 | 447 | 473 | 473 | 594 | 837 | 810 | ID | 972 |
| **N1_1** | 513 | 432 | 54 | 567 | 567 | 54 | 702 | 567 | 567 | 513 | 486 | 729 | 729 | 756 | 756 | 864 | 447 | 447 | 447 | 473 | 594 | 810 | 783 | 972 | ID |

**Figure S2. Comparison of all CRISPR DRs in *Anabaena* 7120. A.** Multiple sequence alignments of all variants of CRISPR DRs in *Anabaena* 7120 and in the cyanophage N-1. See **Figure 1** for the alignment of major variants. The repeats are numbered according to the respective genomic locus as defined previously (Hou et al., 2019). The numbers after the identifier give the copy number of the respective DR (e.g., CR_1 includes 28 copies of the major sequence variant, DR1_28, and one minor variant, DR1_1) followed by a single-letter suffix if needed. **B.** Matrix comparing the repeat identities against each other.

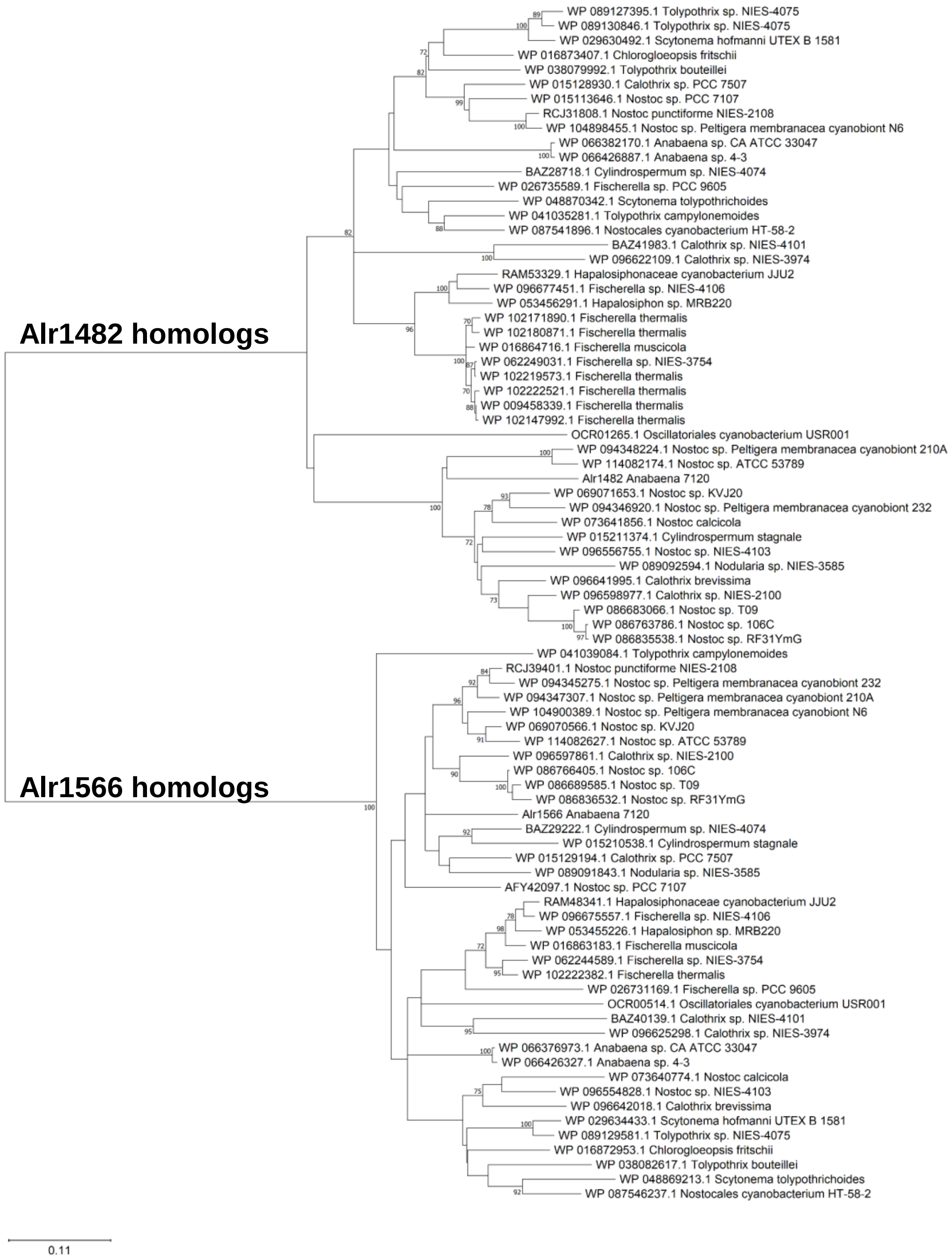


**Figure S3. Phylogenetic analysis of the two Cas6 proteins from *Anabaena* 7120 and their homologs.** The phylogeny was inferred by Minimum Evolution (Rzhetsky and Nei, 1992) as implemented in MEGA X (Kumar et al., 2018). The optimal tree with the sum of branch length = 7.5698 is shown. The percentage of replicate trees in which the associated taxa clustered together in the bootstrap test (1000 replicates) are shown next to the branches if ≥70% (Felsenstein, 1985). The tree is drawn to scale, with branch lengths in the same units as those of the evolutionary distances used to infer the phylogenetic tree. The evolutionary distances were computed using the Poisson correction method (Zuckerkandl and Pauling, 1965) and are in the units of the number of amino acid substitutions per site. The ME tree was searched using the Close-Neighbor-Interchange (CNI) algorithm (Nei and Kumar, 2000) at a search level of 1. The Neighbor-joining algorithm (Saitou and Nei, 1987) was used to generate the initial tree. This analysis involved 82 amino acid sequences. All ambiguous positions were removed for each sequence pair (pairwise deletion option). There were a total of 440 positions in the final dataset.

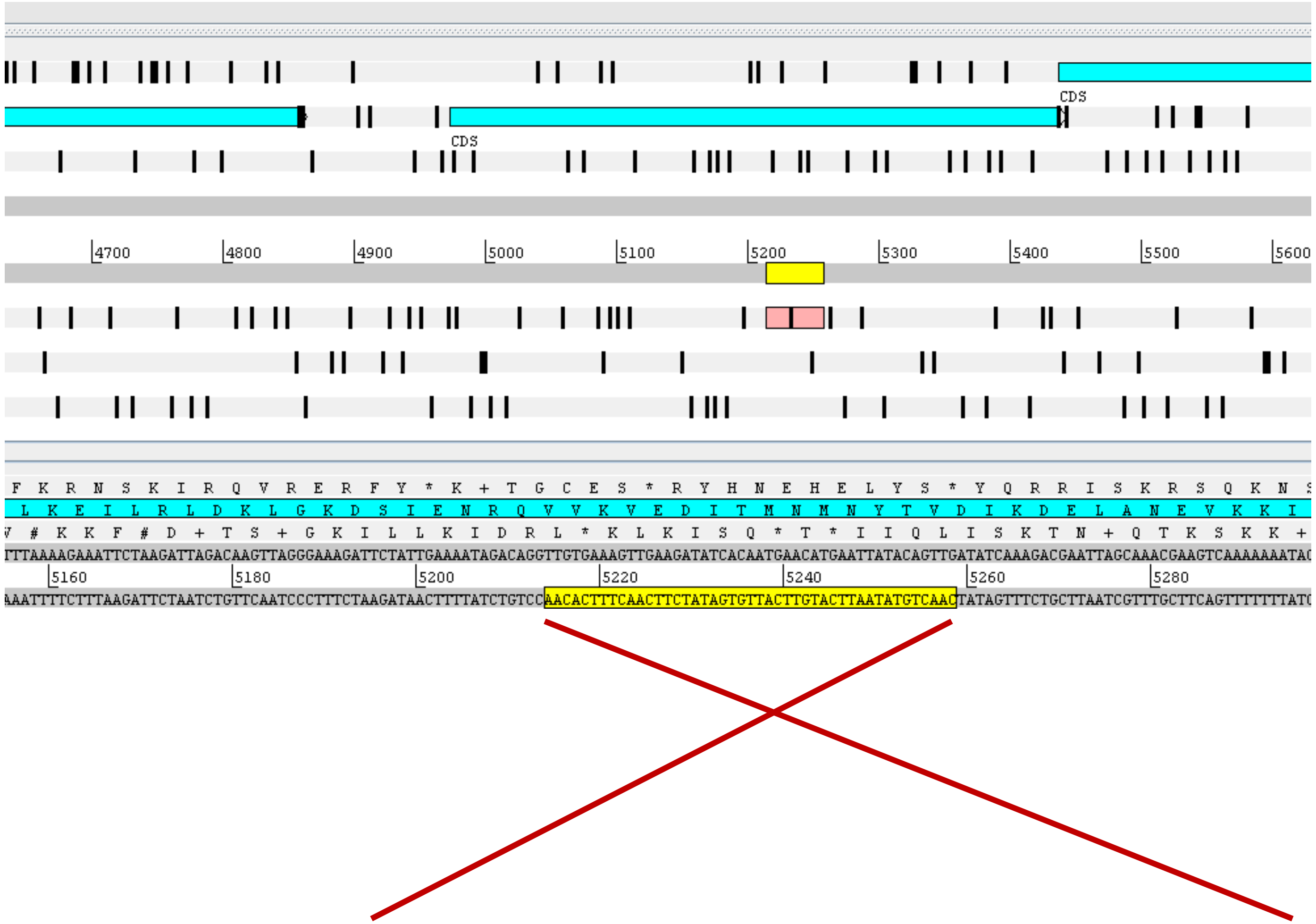


```
N1_c5214..5258: CAACTGTATAATTCATGTTCATTGTGATATCTTCAACTTTCACAA
                 | ||||||||||||| ||||||||||||||||| ||||| ||||
DR1_12:         GACCTGTATAATTCATATTCATTGTGATATCTTCCACTTTTACAA
```

**Figure S4. Targeting of the cyanophage N-1 by the CR_1 system of *Anabaena* 7120.** The upper part shows the genomic region in phage N-1 containing a likely protospacer matching the sequence of spacer 12 of the CR-1 system of *Anabaena* 7120 (lower part).

**Next page:**

**Figure S5. Analysis of *cas6* deletion mutants for the CR_2/3 array *in vivo*.**

**A.** The accumulation of crRNAs and precursors for the split CR_2/3 repeat-spacer array after RNA separation in high-resolution polyacrylamide gels and Northern hybridization. The blots show RNA hybridization results with three clones each from WT and the deletion mutants Δ*alr1482* and Δ*alr1566*. **B.** On top, the coverage in a small RNA transcriptome in WT cells is shown along the full length of the respective array (grey, repeats; yellow, spacers) and below in detail for a selected number of reads. The color of the reads highlights their direction (red: forward; blue: reverse) and mutations inside the reads are marked by different colors (adenine: green; cytosine: blue; guanine: orange; thymine: red). The sequence below represents the DNA sequence of the sense strand of the CRISPR array. The region recognized by the respective probe in the Northern hybridization in panel (A) is indicated by the orange arrow. The orientation of the CRISPR arrays is illustrated by black arrows.

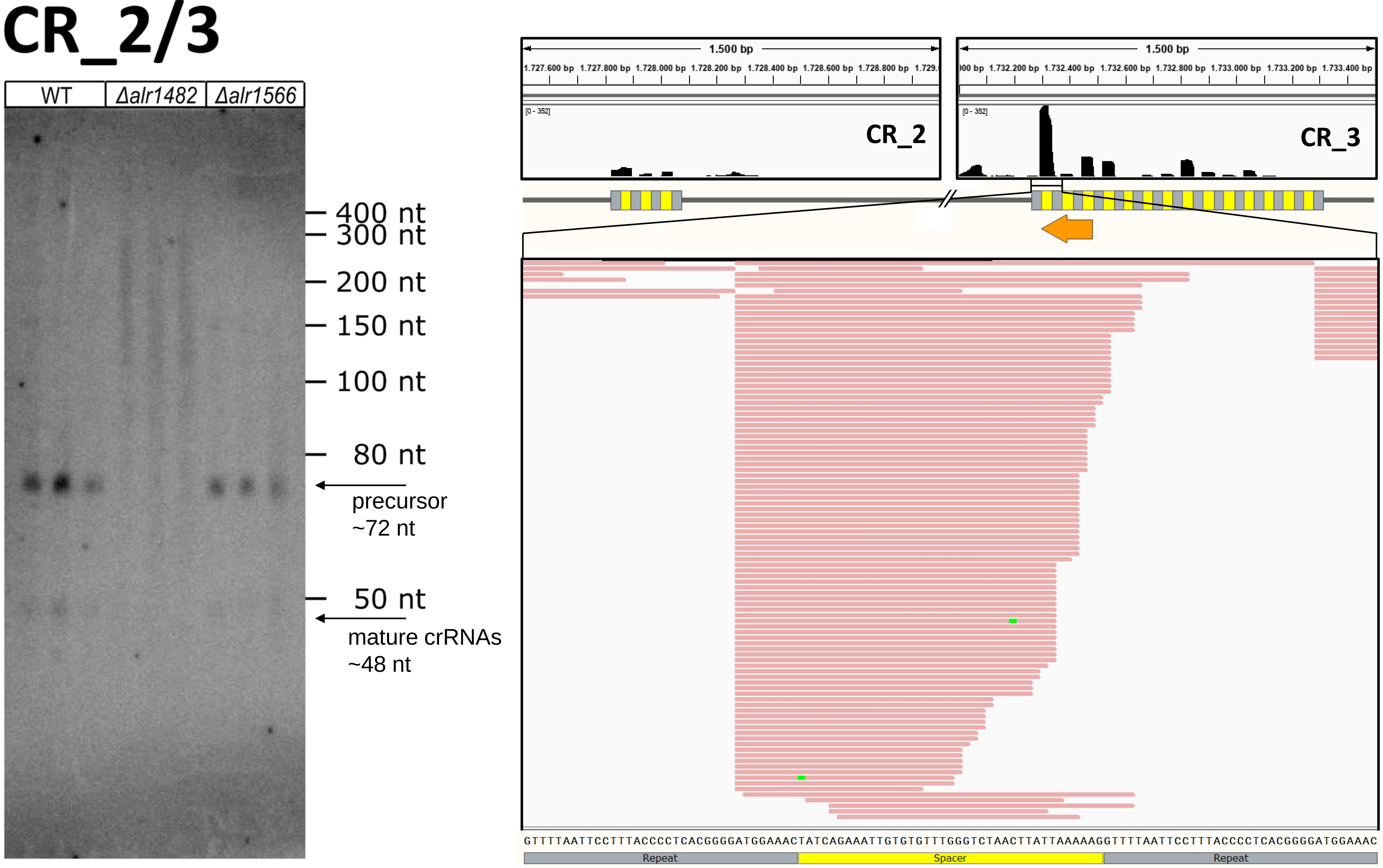
CR_2/3
WT
Δalr1482
Δalr1566
400 nt
300 nt
200 nt
150 nt
100 nt
80 nt
precursor
~72 nt
50 nt
mature crRNAs
~48 nt
1.500 bp
1.727.600 bp
1.727.800 bp
1.728.000 bp
1.728.200 bp
1.728.400 bp
1.728.600 bp
1.728.800 bp
[0 - 352]
CR_2
1.500 bp
1.732.200 bp
1.732.400 bp
1.732.600 bp
1.732.800 bp
1.733.000 bp
1.733.200 bp
1.733.400 bp
[0 - 352]
CR_3
GTTTTAATTCCTTTACCCCTCACGGGGATGGAAACTATCAGAAATTGTGTGTTTGGGTCTAACTTATTAAAAAGGTTTTAATTCCTTTACCCCTCACGGGGATGGAAAC
Repeat
Spacer
Repeat

**Figure S5**

**Next page:**

**Figure S6. Analysis of *cas6* deletion mutants for the CR_5 array *in vivo*.**

**A.** The accumulation of crRNAs and precursors for the CR_5 repeat-spacer array after RNA separation in high-resolution polyacrylamide gels and Northern hybridization. The blots show RNA hybridization results with three clones each from WT and the deletion mutants Δ*alr1482* and Δ*alr1566*. **B.** On top, the coverage in a small RNA transcriptome in WT cells is shown along the full length of the respective array (grey, repeats; yellow, spacers) and below in detail for a selected number of reads. The color of the reads highlights their direction (red: forward; blue: reverse) and mutations inside the reads are marked by different colors (adenine: green; cytosine: blue; guanine: orange; thymine: red). The sequence below represents the DNA sequence of the sense strand of the CRISPR array. The region recognized by the respective probe in the Northern hybridization in panel (A) is indicated by the orange arrow. The orientation of the CRISPR array is illustrated by black arrows.

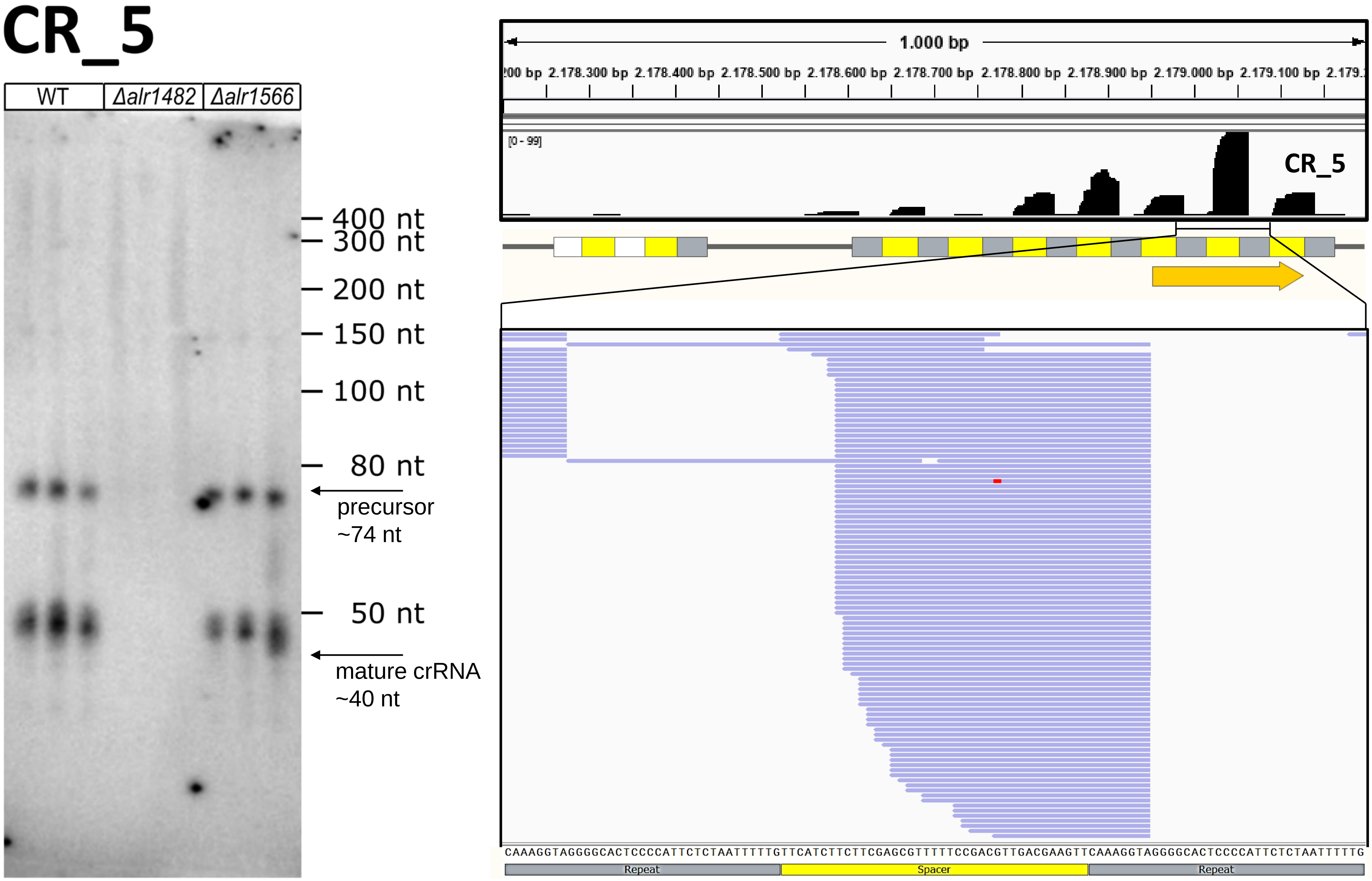
CR_5
WT
Δalr1482
Δalr1566
400 nt
300 nt
200 nt
150 nt
100 nt
80 nt
precursor ~74 nt
50 nt
mature crRNA ~40 nt
1.000 bp
2.178.300 bp 2.178.400 bp 2.178.500 bp 2.178.600 bp 2.178.700 bp 2.178.800 bp 2.178.900 bp 2.179.000 bp 2.179.100 bp
[0 - 99]
CR_5
CAAAGGTAGGGGCACTCCCCATTCTCTAATTTTTGTTCATCTTCTTCGAGCGTTTTTCCGACGTTGACGAAGTTCAAAGGTAGGGGCACTCCCCATTCTCTAATTTTTG
Repeat
Spacer
Repeat

Figure S6

**Next page:**

**Figure S7. Analysis of *cas6* deletion mutants for the CR_10 array *in vivo*.**

**A.** The accumulation of crRNAs and precursors for the CR_10 repeat-spacer array after RNA separation in high-resolution polyacrylamide gels and Northern hybridization. The blots show RNA hybridization results with three clones each from WT and the deletion mutants Δ*alr1482* and Δ*alr1566*. **B.** On top, the coverage in a small RNA transcriptome in WT cells is shown along the full length of the respective array (grey, repeats; yellow, spacers) and below in detail for a selected number of reads. The color of the reads highlights their direction (red: forward; blue: reverse) and mutations inside the reads are marked by different colored inserts (adenine: green; cytosine: blue; guanine: orange; thymine: red). The sequence below represents the DNA sequence of the sense strand of the CRISPR array. The region recognized by the respective probe in the Northern hybridization in panel (A) is indicated by the orange arrow. The orientation of the CRISPR array is illustrated by a black arrow.

# CR_10

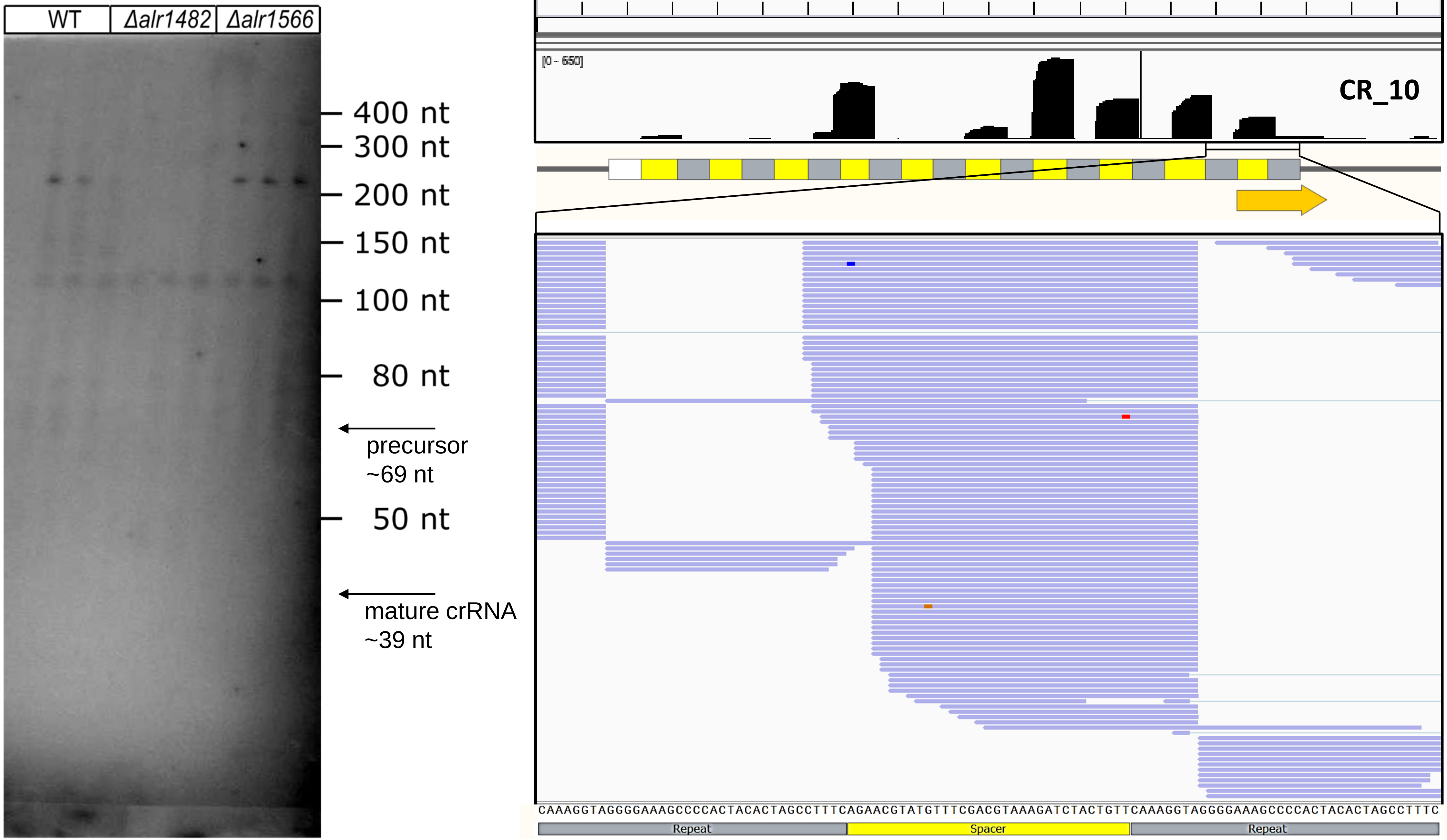


Figure S7

**Next page:**

**Figure S8. Analysis of *cas6* deletion mutants for the CR_4 array *in vivo*.**

**A.** The accumulation of crRNAs and precursors for the CR_4 repeat-spacer array after RNA separation in high-resolution polyacrylamide gels and Northern hybridization. The blots show RNA hybridization results with three clones each from WT and the deletion mutants Δ*alr1482* and Δ*alr1566*. **B.** On top, the coverage in a small RNA transcriptome in WT cells is shown along the full length of the respective array (grey, repeats; yellow, spacers) and below in detail for a selected number of reads. The color of the reads highlights their direction (red: forward; blue: reverse) and mutations inside the reads are marked by different colors (adenine: green; cytosine: blue; guanine: orange; thymine: red). The sequence below represents the DNA sequence of the sense strand of the CRISPR array. The region recognized by the respective probe in the Northern hybridization in panel (A) is indicated by the orange arrow. The orientation of the CRISPR array is illustrated by a black arrow.

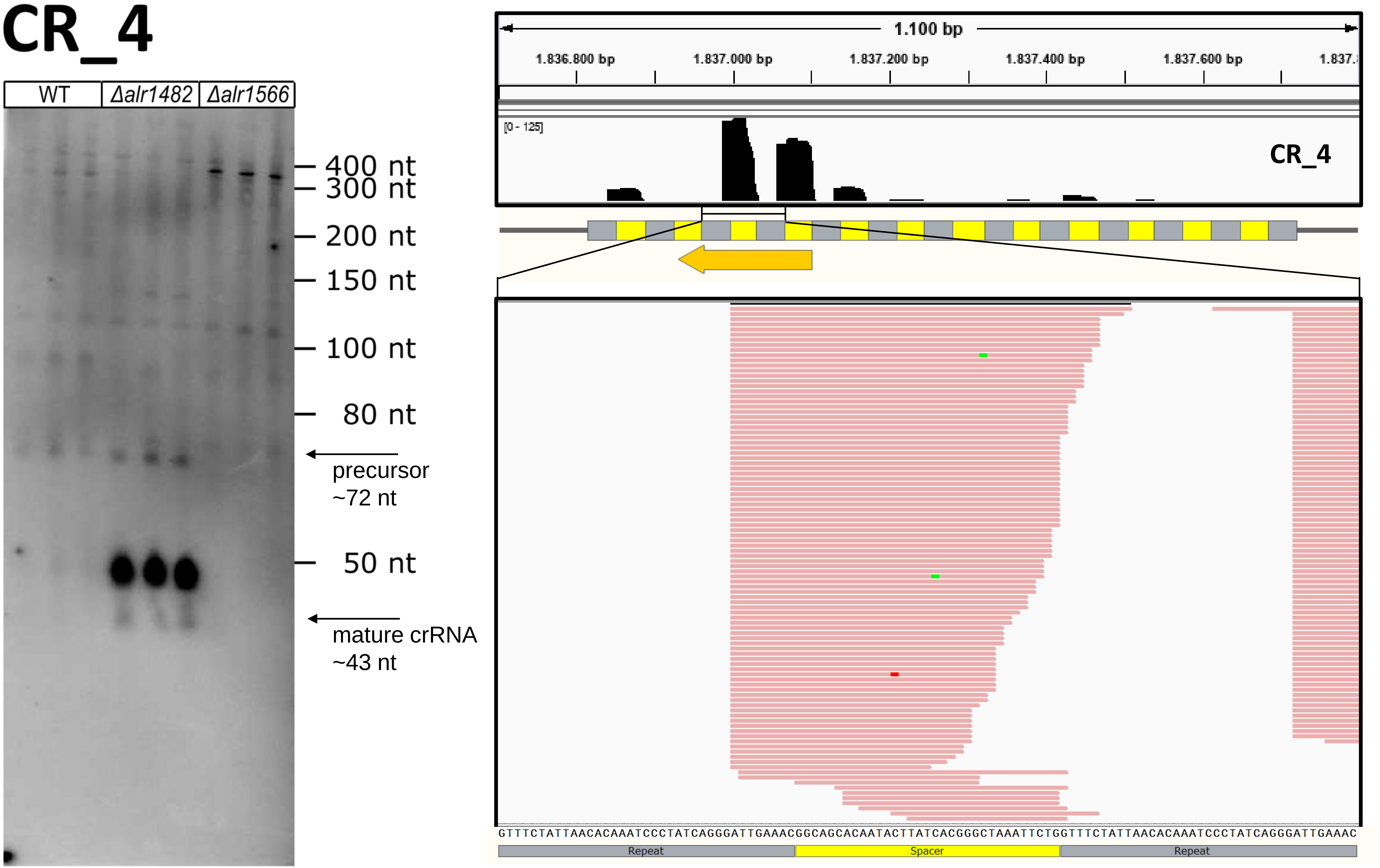

CR_4
WT
Δalr1482
Δalr1566
400 nt
300 nt
200 nt
150 nt
100 nt
80 nt
precursor
~72 nt
50 nt
mature crRNA
~43 nt
1.100 bp
1.836.800 bp
1.837.000 bp
1.837.200 bp
1.837.400 bp
1.837.600 bp
[0 - 125]
CR_4
GTTTCTATTAACACAAATCCCTATCAGGGATTGAAACGGCAGCACAATACTTATCACGGGCTAAATTCTGGTTTCTATTAACACAAATCCCTATCAGGGATTGAAAC
Repeat
Spacer
Repeat


Figure S8

**Next page:**

**Figure S9. Analysis of *cas6* deletion mutants for the CR_6/7 array *in vivo*.**

**A.** The accumulation of crRNAs and precursors for the CR_6/7 repeat-spacer array which is interrupted by a MITE element. RNA was separated in high-resolution polyacrylamide gels and Northern hybridization. The blots show RNA hybridization results with three clones each from WT and the deletion mutants Δ*alr1482* and Δ*alr1566*. **B.** On top, the coverage in a small RNA transcriptome in WT cells is shown along the full length of the respective array (grey, repeats; yellow, spacers) and below in detail for a selected number of reads. The color of the reads highlights their direction (red: forward; blue: reverse) and mutations inside the reads are marked by different colors (adenine: green; cytosine: blue; guanine: orange; thymine: red). The sequence below represents the DNA sequence of the sense strand of the CRISPR array. The region recognized by the respective probe in the Northern hybridization in panel (A) is indicated by the orange arrow. The orientation of the CRISPR arrays is illustrated by black arrows.

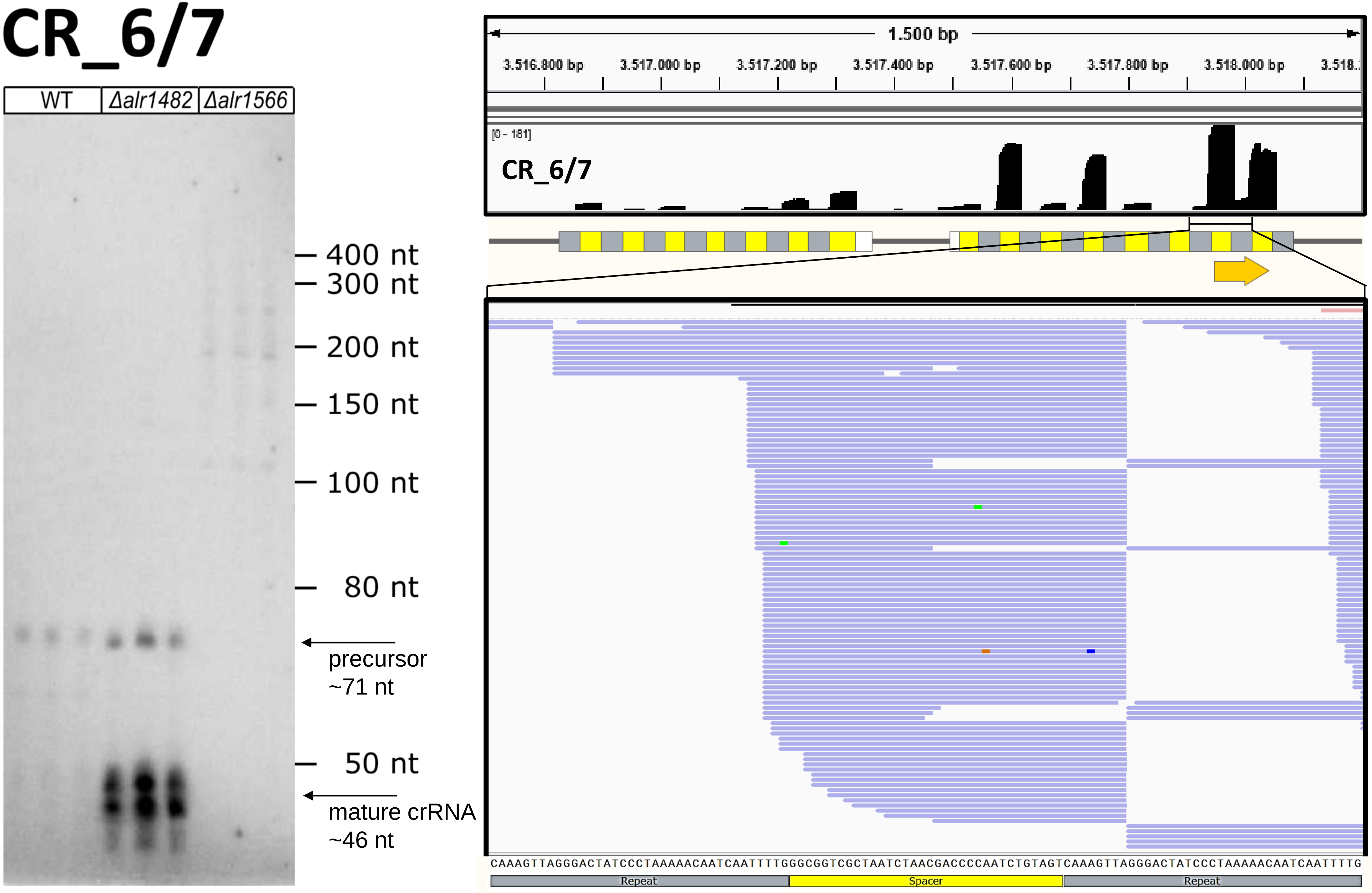
CR_6/7
WT
Δalr1482
Δalr1566
400 nt
300 nt
200 nt
150 nt
100 nt
80 nt
precursor
~71 nt
50 nt
mature crRNA
~46 nt
1.500 bp
3.516.800 bp
3.517.000 bp
3.517.200 bp
3.517.400 bp
3.517.600 bp
3.517.800 bp
3.518.000 bp
[0 - 181]
CR_6/7
CAAAGTTAGGGACTATCCCTAAAAACAATCAATTTTGGGCGGTCGCTAATCTAACGACCCCAATCTGTAGTCAAAGTTAGGGACTATCCCTAAAAACAATCAATTTTG
Repeat
Spacer
Repeat

Figure S9

**Next page:**

**Figure S10. Analysis of *cas6* deletion mutants for the CR_11 array *in vivo*.**

**A.** The accumulation of crRNAs and precursors for the CR_11 repeat-spacer array after RNA separation in high-resolution polyacrylamide gels and Northern hybridization. The blots show RNA hybridization results with three clones each from WT and the deletion mutants Δ*alr1482* and Δ*alr1566*. **B.** On top, the coverage in a small RNA transcriptome in WT cells is shown along the full length of the respective array (grey, repeats; yellow, spacers) and below in detail for a selected number of reads. The color of the reads highlights their direction (red: forward; blue: reverse) and mutations inside the reads are marked by different colors (adenine: green; cytosine: blue; guanine: orange; thymine: red). The sequence below represents the DNA sequence of the sense strand of the CRISPR array. The region recognized by the respective probe in the Northern hybridization in panel (A) is indicated by the orange arrow. The orientation of the CRISPR array is illustrated by a black arrow.

# CR_11

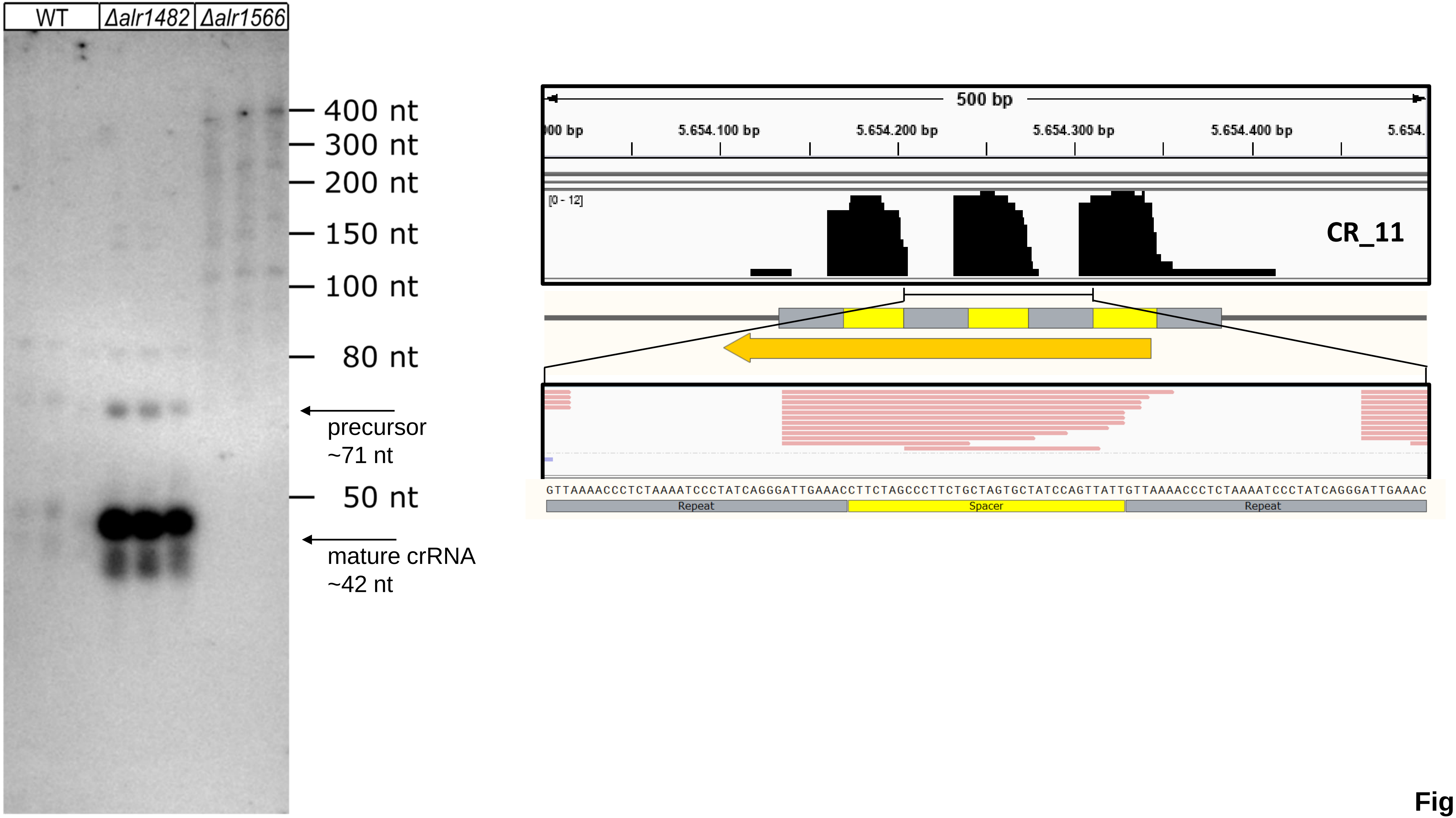


Figure S10
